# Spatially Resolved Reaction–Diffusion Modeling Reveals Effects of Intracellular Spatial Heterogeneity on Yeast Galactose Network Dynamics

**DOI:** 10.1101/2025.07.23.666409

**Authors:** Tianyu Wu, Marie-Christin Spindler, Abner T. Apsley, Emmy Earnest, Zane R. Thornburg, Julia Mahamid, Zaida Luthey-Schulten

## Abstract

Eukaryotic cells are spatially organized into functionally-distinct compartments. This three-dimensional(3D) organization generates intracellular heterogeneities that can modulate regulatory dynamics. Despite this knowledge of subcellular organization, most quantitative gene-regulation models still assume a well-mixed environment in which molecules can react regardless of their spatial positions. Here, we use the well-established galactose switch in budding yeast (*Saccharomyces cerevisiae*) to develop spatially-resolved models that integrate experimentally-derived intracellular architectures, including chromosome organization, the endoplasmic reticulum (ER) and spatially distinct ribosome populations. We implement a hybrid stochastic–deterministic framework in which gene expression is modeled using a reaction–diffusion master equation that enforces locality (i.e., reactions occur only when molecules are in physical proximity), while metabolic and transport processes are captured by ordinary differential equations. Guided by electron microscopy and biochemical constraints, we quantify how accounting for intracellular spatial organization alters regulatory predictions in the galactose switch. We show that chromosome geometry has little effect on Gal2p output, whereas ER-associated translation reduces Gal2p delivery to the plasma membrane; the largest decrease of Gal2p abundance occurs when translation of *GAL2* mRNA is restricted to a population of ribosomes physically bound to the ER. Together, these results demonstrate that more realistic 3D cellular architectures and local reaction rules can qualitatively change regulatory predictions, motivating integration of intracellular organization in future whole-cell models.

**Author Summary:** Eukaryotic cells are highly organized spaces with distinct subcellular compartments and heterogeneous distributions of molecules which influence how cells function. Yet, most computational models of gene regulation assume a spatially homogeneous intracellular environment. To quantify how intracellular architecture influences regulatory predictions, we developed spatially resolved models of the galactose switch in budding yeast, a well-characterized gene regulatory system controlling the response to extracellular galactose. We compared conventional well-stirred simulations with models that explicitly incorporate experimentally informed intracellular organization, including chromosome positioning, endoplasmic reticulum geometry, and functionally distinct ribosome populations. Incorporating these spatial features substantially altered the dynamics of predicted gene activity, protein production, and intracellular sugar levels. Our results demonstrate that the three-dimensional organization of eukaryotic cells can significantly change regulatory outcomes of computational models, underscoring the need for integrating realistic spatial architectures in future models of eukaryotic gene regulation.

## 1 Introduction

Eukaryotic cells are characterized not only by the complexity of their metabolic networks but also by the spatial heterogeneity imposed by their internal organization [1, 2]. Molecules therefore experience non-uniform microenvironments shaped by diffusion constraints, including macromolecular interactions [3] and crowding [4, 5]the positioning of membrane-bound organelles [6]. These spatial features influence the timing and probability of molecular encounters [4, 5], the accessibility of genomic loci [7, 8], and the establishment of compartment-specific biochemical states. As a result, the physical organization of a eukaryotic cell is an active determinant of gene-regulatory dynamics.

Quantitative models that incorporate both biochemical kinetics and spatial constraints are required for understanding how intracellular organization shapes molecular regulation [9]. Whole-cell modeling frameworks provide this capability by integrating reaction kinetics, molecular diffusion, macromolecular interactions, and organelle geometries into a unified simulation environment [10, 11]. When spatial information is explicitly represented, these models can reveal qualitative and quantitative regulatory behaviors that do not arise in well-stirred, homogeneous descriptions [12, 13].

Despite these advantages, building spatially-resolved models of eukaryotic cells remains technically challenging. Most existing whole-cell models focus on prokaryotes, whose smaller volumes and simpler architectures compared to eukaryotic cells reduce the need to account for complex organelle geometries and heterogeneous diffusive environments [14]. Extending these frameworks to eukaryotes requires incorporating genome organization, organelle-specific geometries, heterogeneous transport pathways, and spatially regulated transcription, RNA transport, translation and protein localization [15–17]. A practical strategy for modeling eukaryotic cells is to first construct spatially-resolved models of well-characterized molecular subsystems and incrementally introduce additional layers of biological complexity. In this context, complexity refers specifically to 1) diffusion-induced differences across the cell, where finite mixing times in eukaryotic-scale volumes generate concentration gradients and locally distinct biochemical environments and 2) the explicit representation of major organelles and their associated biochemical processes. Subsystem-level models offer a tractable platform for testing how spatial features influence regulatory behaviors before scaling up to full cellular complexity.

The galactose switch in budding-yeast (*Saccharomyces cerevisiae*) is an ideal molecular subsystem for investigating how spatial heterogeneity modifies regulatory dynamics [18]. Yeast must selectively express galactose-related (*GAL*) genes when galactose is available as the primary carbon source, and this regulatory decision requires coordinated interactions among extracellular sugar, sensors, gene repressors and activators, and transmembrane transporters. Although analogous regulatory switches exist in bacteria [19–21], the larger size and compartmental complexity of eukaryotic cells create spatial environments in which diffusion, organelle proximity, and intracellular trafficking can significantly influence the kinetics of regulatory interactions. These differences make the galactose switch a powerful test-bed for assessing how spatial context governs regulatory outcomes.

In budding yeast in particular, a wealth of ultrastructural and structural information is available across diverse imaging modalities [22–27]. Whole-cell and organelle-scale 3D reconstructions derived from volumetric imaging including electron tomography of resin-embedded samples, soft X-ray tomography, and focused ion beam-scanning electron microscopy (FIB-SEM) volume imaging have revealed the 3D organization of the intracellular architecture at nanometer to tens-of-nanometer resolution. These datasets provide direct geometric constraints on organelle morphology, spatial relationships, and relative positioning of organelles within intact cells [24–27]. Quantitative cryo-electron tomography (cryo-ET) studies further provide measurements of macromolecular state, concentrations, organelle association and spatial organization across growth conditions and cell cycle states, making yeast an unusually well-characterized eukaryotic system for deriving spatial priors used in spatially resolved simulations [28–33]. Several features make the galactose switch particularly suitable for spatially-resolved modeling. First, the genetic and biochemical regulation of the switch have been extensively quantified over decades of study [18, 34]. Second, a well-validated spatially homogeneous stochastic-deterministic hybrid model provides a baseline for evaluating how spatial extensions alter predicted regulatory behavior [35]. Third, galactose-related protein abundances are experimentally accessible [36, 37], enabling direct comparison of experimental observations with simulation results. Together, these features allow systematic evaluation of how spatial heterogeneity reshapes dynamics in a canonical eukaryotic regulatory circuit.

Here, we report the construction of a spatially resolved model of the yeast galactose switch to evaluate how realistic subcellular architectures influence regulatory behavior (Figure 1). To capture both spatially localized reactions and global biochemical processes, we implemented a hybrid stochastic–deterministic framework in which reaction–diffusion dynamics were simulated using a multi-GPU RDME–ODE solver in Lattice Microbes [38]. This implementation enabled efficient large-scale simulations through hardware-aware optimizations, including CPU–GPU affinity mapping, reducing runtime from approximately 10 hours to 6 hours.

**Figure 1.**
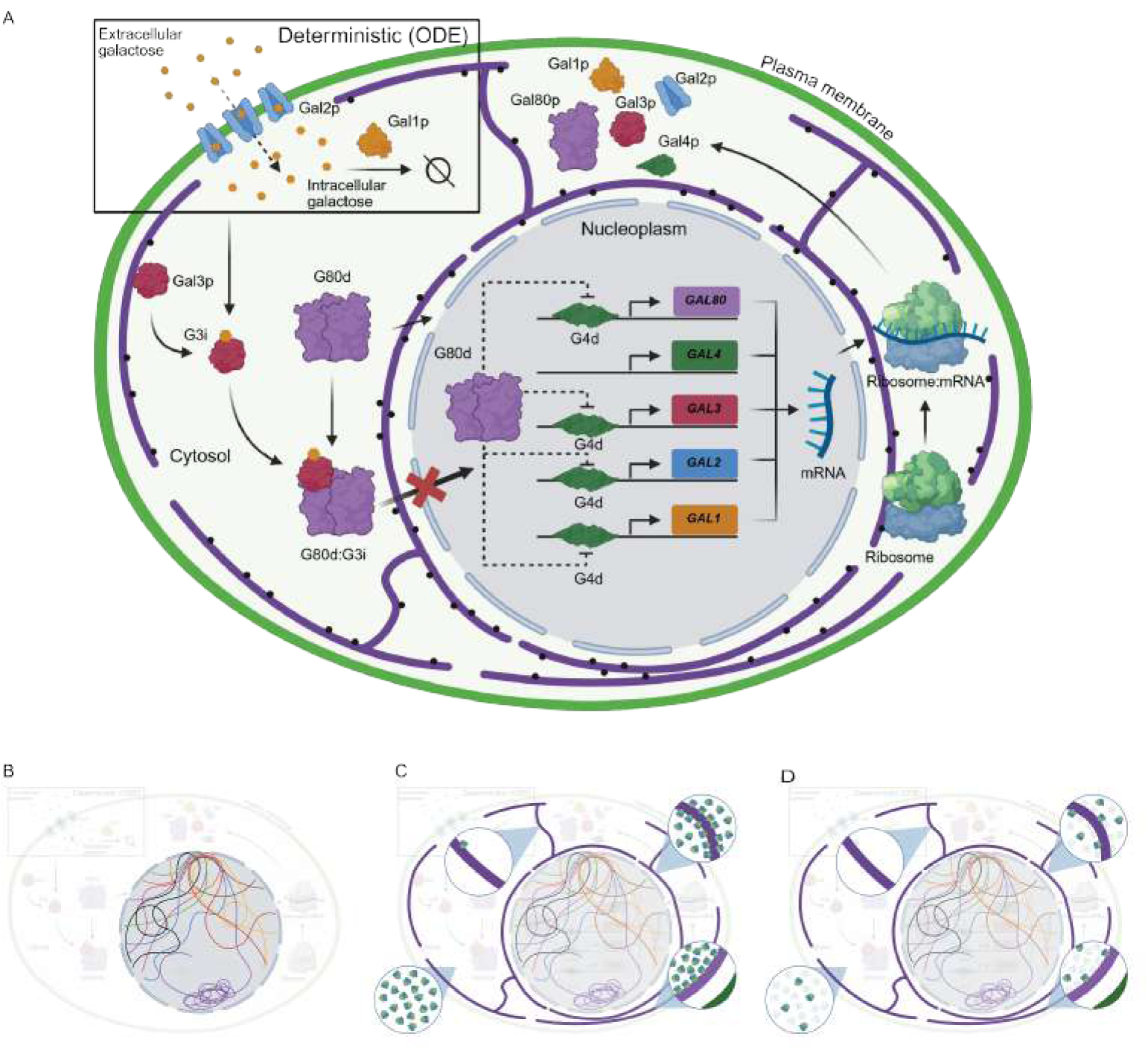
Yeast galactose switch system and study design. (A) Schematic of the yeast galactose switch system. G4d activates *GAL* gene expression, whereas G80d represses it.. The suffix “d” indicates the dimeric state and “p” indicates the monomeric protein. Black dots on the ER denote ER-associated ribosomes. (B-D) Schematics of the stepwise extension of the spatially-resolved model. (B) Addition of spatial information about the yeast genome. (C) Addition of ER geometry and ER-attached ribosomes. (D) Consideration of decreased number of ribosomes available for the galactose switch system due to translation competition.

Using this framework, we first compared the spatially resolved model with the established well-stirred formulation to isolate regulatory changes arising solely from the introduction of spatial heterogeneity. We then assessed how progressively incorporating additional architectural regulatory features—chromosome organization, ER geometry, ER-associated ribosomes, and system-specific effective ribosome abundance—modulated the switch’s dynamic response. Collectively, these analyses establish a spatially constrained modeling strategy for testing how organelle-scale geometry and localized translation reshape regulatory dynamics, effects that cannot be captured by conventional well-mixed models.

## 2 Materials and Methods

### 2.1 Chemical Kinetics of the Yeast Galactose Switch

#### 2.1.1 Overview of the Yeast Galactose Switch

The yeast galactose switch governs the expression of genes needed for galactose utilization when glucose, the preferred carbon source, is depleted. Our representation of the network comprised 39 molecular species (see Supplementary File 1: Tables S1-S4) and 75 reactions (see Supplementary File 1: Tables S5-S10) arranged in two positive and one negative feedback loops that respond dynamically to the presence of extracellular galactose [18, 34]. As illustrated in Figure 1, four core genes—*GAL1*, *GAL2*, *GAL3*, and the repressor *GAL80* are induced upon encountering intracellular galactose. *GAL4*, also part of the galactose switch system, acts as a constitutively expressed transcriptional activator of all other *GAL* genes. Both Gal4p (*GAL4* protein product) and Gal80p (*GAL80* protein product) exist as either monomers or dimers, but only exert their respective functions on the galactose switch system in the dimeric form. The Gal4p dimer (G4d) binds to a conserved nucleotide sequence called the upstream activation site (UAS; 5’-CGG-N_11_-CCG-3’) which is located upstream of *GAL1*, *GAL2*, *GAL3*, and *GAL80* (see Supplementary File 1: Table S1). In the presence of intracellular galactose, the *GAL3* protein (Gal3p) binds the sugar (forming G3i, the sugar induced form of Gal3p) and prevents the repressor G80d from diffusing into the nucleus, sustaining *GAL* gene expression. The *GAL1* protein (Gal1p) catalyzes the addition of a phosphate group to galactose, the first metabolic step in the Leloire pathway, whereas *GAL2* protein (Gal2p), which is translated at the ER, is shuttled to the plasma membrane where it acts as a galactose transporter. For a complete description of the chemical species present and the reaction network used, see Tables S1-S10.

#### 2.1.2 Simulating Spatially-Resolved Stochastic Chemical Kinetics Using the Reaction-Diffusion Master Equation

The reaction-diffusion master equation (RDME) provides a framework for performing spatially-resolved stochastic simulations of intracellular processes, combining particle-level stochastic reactions with spatial diffusion dynamics. Here, we applied the RDME approach to simulate transcription, translation, and regulator-promoter interactions in the galactose switch. These simulations were performed using the Lattice Microbes (LM) software [38, 39], which supports both well-stirred and spatially-resolved models. For RDME/spatially-resolved models, LM explicitly incorporates spatial effects by discretizing the cellular volume into subvolumes *v* centered on a cubic lattice grid. Reactions occurring within each subvolume are assumed to be well-stirred and are simulated stochastically using the chemical master equation (CME). Each subvolume’s CME is then solved using the Gillespie algorithm [40], while particle diffusion between adjacent subvolumes is modeled using a diffusion operator (see Equation 1).

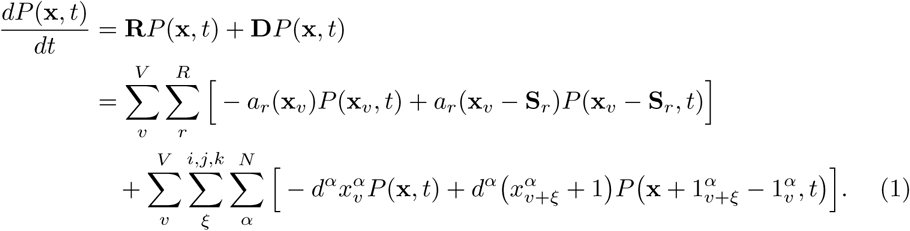

In Equation 1, *P* (**x***, t*) denotes the time-dependent joint probability distribution, representing the likelihood that the system exists in configuration **x** at time *t*. The term *a_r_*(**x***_v_*) is the reaction propensity function, defined such that *a_r_*(**x***_v_*)*dt* represents the probability of reaction *r* occurring in subvolume *v* during the infinitesimal time interval [*t, t* + *dt*), given the current local molecular counts. Consequently, the equation describes the time evolution of *P* (**x***, t*) as a net probability flux—the balance of transitions entering and leaving state **x**—driven by these reaction and diffusion events. The RDME governing stochastic dynamics models the system as a sum of local CMEs within discretized subvolumes and a diffusion operator accounting for transport between them. Specifically, the local CME governs stochastic reactions within each subvolume *v*, while diffusion terms describe the movement of species *α* along spatial directions. The system state **x** encapsulates the discrete molecular counts of all species across all subvolumes.

#### 2.1.3 Modeling Stochastic Processes in the Yeast Galactose Switch Using an RDME Framework

In our present model, genetic information processing reactions (i.e., regulator-promoter interactions, transcription, translation, mRNA degradation, protein degradation, protein complex formation and disassembly) are represented as stochastic processes using an RDME framework (see Supplementary File 1: Tables S5-S10). RDME simulations of the genetic information processing reactions were resolved with subvolume precision, meaning that reactions were only allowed to take place within a specific subcellular location (see Supplementary File 1: Tables S5-S10, “Region” variable). Diffusion behaviors are specified for each molecular species including both diffusion within a given region type and diffusion across boundaries between different regions (see Supplementary File 1: Table S12).

The regulator-promoter interaction events are defined to occur within the nucleoplasm region for GAL genes *GAL1*, *GAL2*, *GAL3*, repressor gene *GAL80*, and the activator gene *GAL4* (see Supplementary File 1: Table S5). For the regulated GAL genes, each has 3 states: free, activated and repressed. When genes are in activated states, transcription is enabled and results in the release of the messenger RNAs (mRNAs) in the same sublattice volume. For the non-regulated gene *GAL4*, we modeled the transcription to take place stochastically with a given rate (see Supplementary File 1: Table S6). The messengers then diffuse from the nucleoplasm to the cytosol through nuclear pores. In the cytosol, mRNAs are either degraded (see Supplementary File 1: Table S7) or diffuse to and bind to ribosomes for translation (see Supplementary File 1: Table S8).

We allow protein complexes to form from three individual protein subunits: Gal3p, Gal4p and Gal80p (see Supplementary File 1: Table S9). First, Galp4 and Gal80p are able to bind with themselves to form the homodimers of G4d and G80d, respectively. Second, the Gal3p subunit is allowed to bind to cytosolic galactose to form a complex G3i, which can then further bind to the G80d in the cytosol forming the G3iG80d complex. Formation of the G3iG80d complex prevents G80d from transporting through the nuclear pore into the nucleoplasm, thereby inhibiting the repressive effects of G80d. Third, G4d–G80d binding to repress gene expression is allowed but only in the nucleoplasm and in the presence of DNA. We adopted this strategy because the C-terminal activation domain (AD; residues 840–881) of G4d is intrinsically disordered in solution, but adopts a structured conformation upon binding to DNA and then G80d — consistent with partner-induced folding typical of intrinsically disordered activation domains [41]. This mechanistic insight justified modeling G4d–G80d complex formation as a nuclear, DNA-dependent event.

#### 2.1.4 Modeling Deterministic Processes in the Yeast Galactose Switch Using an Ordinary Differential Equation Framework

Both the transport of extracellular galactose into the yeast cell via Gal2p and galactose metabolism via Gal1p are modeled deterministically using ordinary differential equations (ODEs; see Table S11). ODEs are solved numerically using the LSODA function in the SciPy package in Python [42]. As our present model does not explicitly account for downstream metabolic pathways, the metabolic product galactose-1-phosphate is thus represented as a dead-end.

#### 2.1.5 Combining Stochastic and Deterministic Reactions into a Hybrid RDME-ODE Method

We employ a hybrid RDME-ODE framework to efficiently simulate spatially-resolved gene expression and metabolism in budding yeast, using stochastic methods for low-copy processes and ODEs for high-copy metabolic reactions. Communication between the RDME and ODE solvers occur at regular intervals *τ* = 1*s* as depicted in Figure 2. The communication time is determined based on the separation of timescales between slow stochastic reactions and faster deterministic metabolic processes [35]. At each communication step, the state of the RDME system (e.g., local particle counts) is updated to reflect the results of the ODE solver. After this, the ODE solver is run and subsequent solutions are communicated back to the RDME framework, thereby updating the RDME kinetic parameters. A description of this algorithm can be found in Algorithm 1.

**Figure 2.**
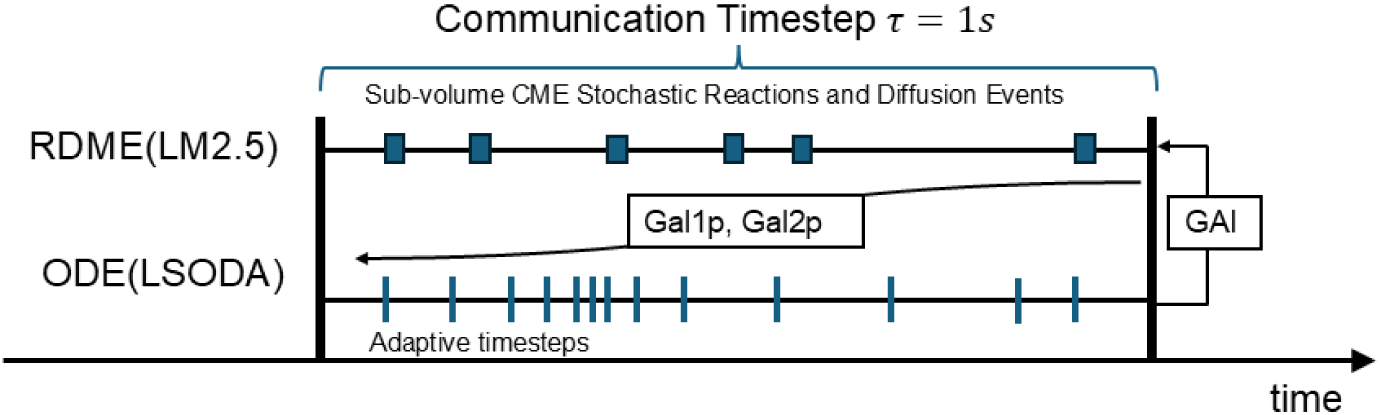
Schematics for the communication procedure between the RDME and ODE submodels. Stochastic processes are modeled using the RDME submodel and deterministic processes are modeled using the ODE submodel. After each RDME step, information about Gal1p and Gal2p abundances are then communicated to the ODE submodel. After each adaptive step in the ODE submodel, intracellular galactose concentrations are communicated back to the RDME submodel.

Notably, the bimolecular reaction *G*3 + *GAI* → *G*3*i* (where *GAI* represents intracellular galactose), characterized by a low copy number of Gal3p and a high copy number of intracellular galactose, posed computational challenges when directly included in the RDME framework. Direct incorporation significantly slowed down the RDME simulations and led to spatially crowded subvoxels due to excessive galactose molecules. To address this, we converted the bimolecular reaction into a pseudo-first-order reaction *G*3 → *G*3*i* by redefining its kinetic parameter as *k^′^* = *k* · [GAI], where *k* represents the original kinetic rate constant and [GAI] denotes the intracellular concentration of galactose. This modification maintains biological accuracy while significantly enhancing computational efficiency. This conversion and parameter update are handled dynamically by the *setReactionRate* function within the simulation hook, newly implemented here for compatibility with multi-GPU solvers.

#### 2.1.6 Defining Initial Abundances of Chemical Species

We define the initial abundances of all chemical species as the values reported in ODE model of Ramsey *et al.* [18], rounding each to the nearest integer (see Tables S2 and S3). For molecular species that are present in multiple cellular compartments, we distribute their initial counts proportionally according to regional volumes. The initial counts used in the RDME simulations are summarized in Tables S2 and S3. For the initial RDME-ODE comparison, *GAL* gene copy numbers are set to one and placed randomly in the nucleoplasm in their repressed state, consistent with biological expectations in the absence of induction. Subsequently, genes are placed in their respective location according to the static chromosome locations included. In both cases, these repressed genes are assumed to be immobile (non-diffusive) throughout the simulation. In contrast, *GAL4*, which is not under regulatory control in the model, is constitutively transcribed. All mRNAs are initialized by rounding the initial abundance used by Ramsey et al. [18](Table S2). The protein species Gal1p and Gal3p are randomly distributed in the cytosol at initialization, while Gal2p is localized to the plasma membrane. G4d is initialized in the nucleoplasm. Gal80p and G80d are randomly distributed between the cytosol and the nucleoplasm in proportion with the volume ratio of the two regions. The initial intracellular galactose concentration is set to zero to reflect experimental conditions in which cells were pre-cultured in a non-inducing medium (raffinose) prior to galactose exposure. In our earlier CME–ODE study [35], quantitative comparisons to the Ramsey ODE model were limited to lower external galactose concentrations due to computational constraints associated with stochastic–deterministic coupling; under these conditions, approximating Hill-type regulation with first- and second-order mass-action reactions yielded Gal2p levels within 5% the Ramsey model predictions [18]. In the present work, extracellular galactose is fixed at 11.1 mM, matching the concentration used in Ramsey et al. [18], to ensure consistency across experimental benchmarks. Simulations are initialized from a repressed transcriptional state with zero intracellular galactose, reflecting the experimental induction protocol in which cells were shifted from raffinose to galactose [18].

##### Algorithm 1 Hybrid RDME-ODE Simulation of the Galactose Switch Using Lattice Microbes

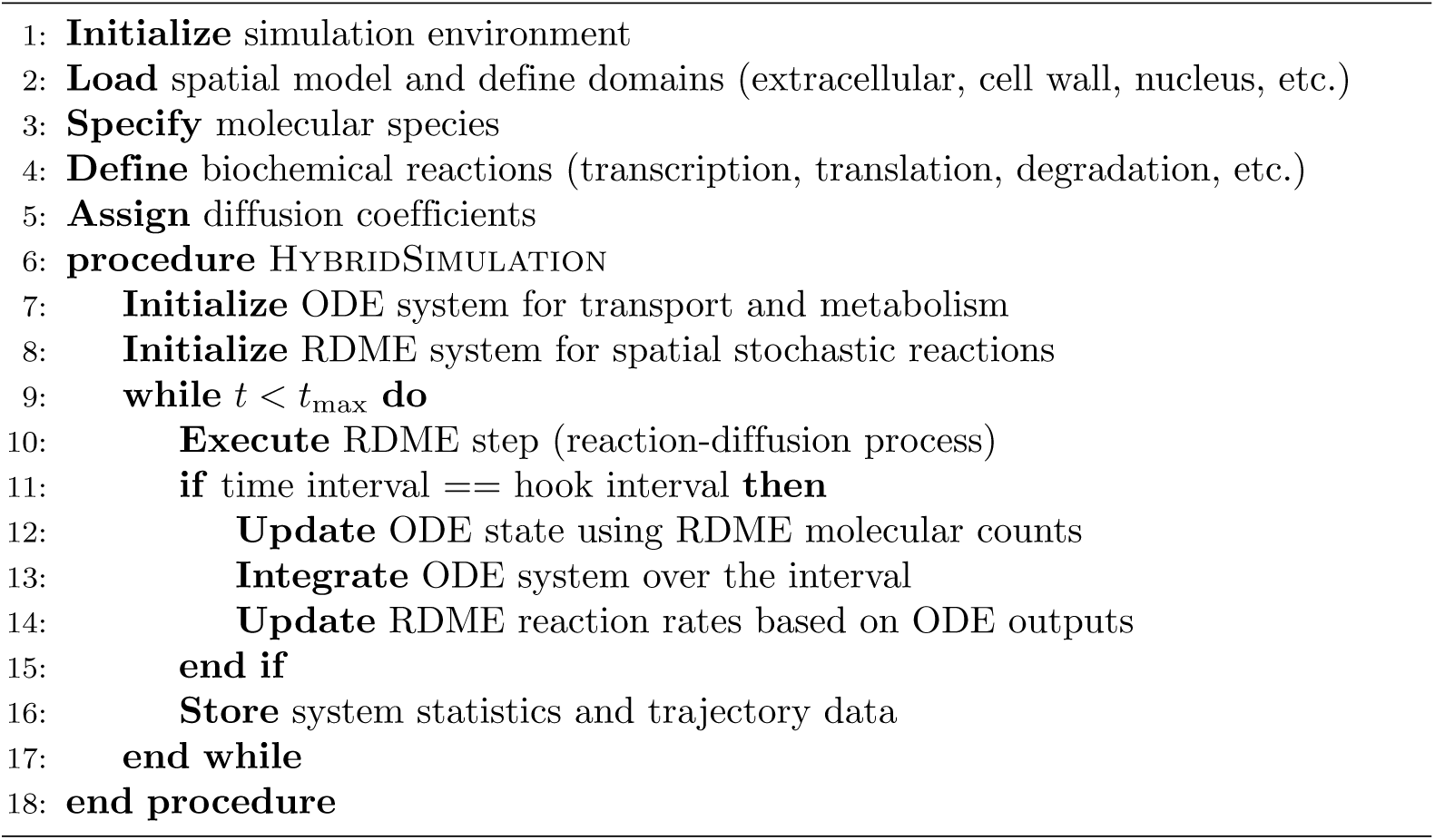

### 2.2 Reconstruction of Yeast Cell Geometry Using Cryogenic Electron Tomography

As described in Earnest et al. [15], the initial 3D geometry of the yeast cell model was built based on a cryo-electron tomogram of a cryo-FIB-milled *S. cerevisiae* cell grown in glucose rich conditions. The tomogram contains a volume of 2.04 × 1.91 × 0.242 *µ*m^3^ and was acquired at a pixel size of 5.31 Å [15]. Due to the limited field of view of the tomogram, missing structures outside the imaged cellular volume were extrapolated based on known volumetric and structural constraints. The final reconstructed geometry was down sampled to a lattice resolution of 28.67 nm to balance simulation accuracy and computational efficiency.

#### 2.2.1 Extrapolation of Ellipsoidal Subcellular Structures

Incomplete geometries in the cryo-ET volume (e.g., partial plasma membrane and nuclear envelope) were completed using the ellipsoid-fitting and extrapolation framework of Earnest et al. [15]. Briefly, segmented membranes were first converted to binary masks (Fig. S1A), and best-fit ellipsoids were obtained by optimizing the nine ellipsoid parameters (three radii, three rotations, and three center coordinates) to match the partial masks. The resulting ellipsoids define the plasma membrane, nuclear envelope, cytosol, and nucleoplasm, and nuclear pores were added following Earnest et al. [15] using the reported pore density (139 pores; shown as white crosses in Fig. S1C).

Additional organelles and structures not fully captured by cryo-ET (mitochondria, vacuole, and cell wall) were reconstructed using the same framework [15], with target volume fractions taken from whole-cell FIB-SEM segmentation measurements [25]. Specifically, the cell wall was modeled as an ellipsoidal shell concentric with the plasma membrane (same center and orientation), with radii chosen to achieve a 15.9% volume occupancy. The vacuole (7.8%) was modeled as a single ellipsoid placed in the cytosol with randomized orientation and positioning constrained to avoid overlap with the nucleus (with semi-axes sampled as in Earnest et al. [15]). Mitochondria (1.6%) were represented as three non-overlapping ellipsoids whose total volume was divided equally among compartments, consistent with BioNumbers (ID:103068) [43, 44]; this simplified representation was used because mitochondria were not explicitly involved in the galactose switch behavior in our model [45] (Fig. 3).

**Figure 3.**
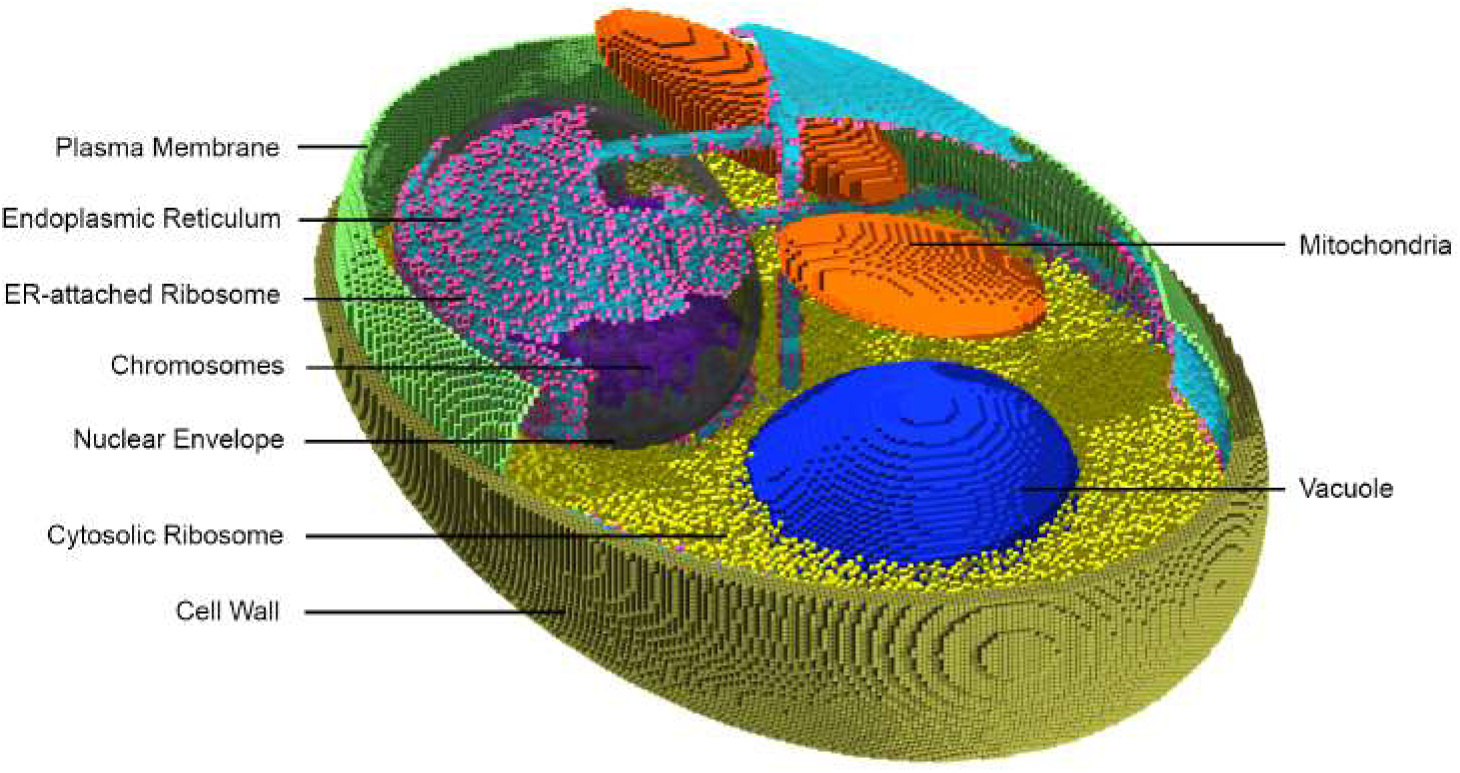
Extended 3D spatial geometry of the model yeast cell. To enhance clarity, only a subset of cytosolic ribosomes, a portion of the cell wall and plasma membrane are shown. From top left to bottom right, the cellular components are colored as follows: plasma membrane (green), endoplasmic reticulum (cyan), ER-attached ribosomes (pink), chromosomes (purple), nuclear envelope (gray), cytosolic ribosomes (yellow), cell wall (tan), mitochondria (orange) and vacuole (blue).

#### 2.2.2 Inclusion of Static Chromosome Geometries

To examine how chromosomal positioning and gene loci influence the galactose switch dynamics, we utilized the static interphase chromosome model of *S. cerevisiae* described by Duan et al. [46]. The model was geometrically transformed and rescaled to maximize spatial occupancy within the nucleoplasm while avoiding intersection with the nuclear envelope. This scaling approach is supported by structural evidence showing that, during interphase, centromeres cluster near one nuclear pole, where they remain connected to the spindle pole body which is embedded in the nuclear envelope [47, 48]. Telomeres are similarly anchored close to the nuclear periphery [49]. The nucleolus, formed partially by extended rDNA repeats on chromosome XII, localizes to the opposite side of the nucleus and is also in contact with the nuclear envelope [50, 51]. Together, these spatial constraints justify scaling the interphase chromosome model to maximize its spatial occupancy within the nucleoplasm.

#### 2.2.3 Modeling the Distribution of Cytosolic Ribosomes

The dimensions of cytosolic ribosomes in *S. cerevisiae* are approximately 25.4 nm × 27.8 nm × 26.7 nm [52]. Given that the lattice spacing in our RDME model was 28 nm × 28 nm × 28 nm, each ribosome was represented as occupying a single lattice voxel. In the model configuration without endoplasmic reticulum (ER) compartments (see Section 2.4), a total of 345,964 ribosomes were included. This number was extrapolated from an average ribosome density of 14,500 ribosomes per *µ*m^3^ reported in a recent cryoET study on *S. cerevisiae* grown in glucose rich conditions [33]. Ribosomes were initialized by uniformly distributing them throughout the cytosolic space using random selection of lattice sites not occupied by existing intracellular structures. The total ribosome count used in our simulations lies within the experimentally reported range of 105,000–380,414 ribosomes per yeast cell [53–55]. During the first 60 minutes of simulation, the total number of actively translating ribosomes for mRNAs for all *GAL* genes remained below 150 (see Figure S9).

#### 2.2.4 Modeling the Geometries of the Endoplasmic reticulum

The yeast ER network structure differs significantly from the tubular network structures typically observed in mammalian and plant cells [56, 57]. The yeast ER primarily consists of large cisternae near the plasma membrane called the plasma-membrane associated ER (pmaER) and those surrounding portions of the nuclear envelope called cisternal ER (cecER) [58, 59]. The pmaER and cecER are connected by sparse rod-like structures called tubular ER (tubER) [58, 59].

We developed a method based on available electron and fluorescence microscopy data to geometrically model subsections of the ER. First, we used previously reported data, which estimated that the volumetric occupancy of the entire yeast ER was around 2.2% of the total cell volume [25] to specify the volume percent of total ER in our model. We also decomposed the ER network into the three previously described ER subregions: pmaER (50% of the total ER volume), tubER (30% of the total ER volume), and cecER (20% of the total ER volume) [24]. Average thicknesses of the pmaER and cecER sheets were chosen to be 35.6 nm and 36.0 nm, respectively, with a 33.0 nm gap between the ER sheets and the adjacent membranes [24].

To construct the pmaER, we first defined an ellipsoidal shell with a thickness of one lattice cube. This was obtained by subtracting an inner ellipsoid, reduced by two lattice cubes in each semi-axis, from an outer ellipsoid which was itself reduced by one lattice cube in each semi-axis from the plasma membrane. The following equations represent this process:

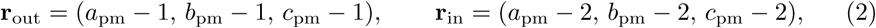

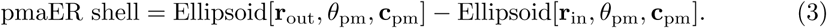

Here, *a*_pm_*, b*_pm_*, andc*_pm_ denote the semi-axes of the plasma membrane ellipsoid, *θ*_pm_ = (*α, β, γ*) represents the pmaER orientation (Euler angles), and **c**_pm_ represents the pmaER’s center. The operator Ellipsoid[**r***, θ,* **c**] represents an ellipsoid with semi-axes **r**, orientation *θ*, and center **c**.

After defining the ellipsoidal shell for the pmaER, we generated pmaER segments by randomly defining *n* fragments with individual volumes {*V_i_* : *i* ∈ [1*, n*]} within the pmaER shell. These fragment volumes were subject to the following constraint:

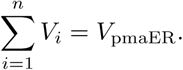

A breadth-first search (BFS) algorithm was applied to iteratively increase the volume of each fragment until the target volume *V_i_* was reached. This process was repeated *n* times, ensuring that no two pmaER fragments overlapped. This procedure is described mathematically below:

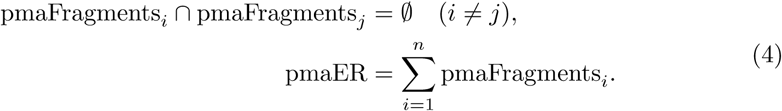

A similar procedure was used to construct the cecER.

In contrast, the tubER was modeled as tubular structures with cross-sectional diameters *d* randomly selected from the range 60–100 nm [60]. The total tubER length was calculated as 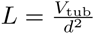. Tubules were generated by connecting randomly selected pairs of pmaER and cecER fragments until the available length was exhausted.Because the ER geometry is static over the course of the simulation, all cecER components were connected to the pmaER via tubER to ensure global ER connectivity. The resulting ER architecture is shown in Supplementary Video 1:.

#### 2.2.5 Modeling ER-Associated Ribosomes

To parametrize translation by ER-bound, translocation-competent ribosomes, that are required for synthesis of membrane proteins (i.e., plasma membrane-localized Gal2p), we determined the densities of these ribosomes on ER membranes based on cryo-electron tomograms of *S. cerevisiae*. To generate these data, cells were grown in complete and synthetic media supplemented with 20 mM dextrose, vitrified by plunge freezing, thinned by cryo-FIB milling and imaged by cryo-ET. A set of 9 cryo-electron tomograms that contains ER regions were reconstructed in IMOD [61] at a pixel size of 13.48 Å. The dense globular ribosomes were clearly discernible in these tomograms, enabling their localization using a pre-trained 3D Convolutional Neuronal Network (CNN) implemented in DeePiCt [62]. The CNN prediction output was manually curated in EMAN2 [63] to achieve roughly a 95 % coverage of ribosome annotations in the data. Based on all detected 80S ribosomes, we generated a subtomogram average. Multiple rounds of 3D classification and refinement in RELION [64] resulted in a consensus map from 10,723 ribosomes with a resolution of 11.1 Å. Based on this average, we determined the position of the peptide exit tunnel in ChimeraX [65]. MemBrain seg [66] was used to automatically annotate all membranes in the tomograms, the ‘find connected regions’ Image J plugin [67] was used to assign labels to the different organelle membranes and to export the binary masks of the ER membranes. Napari [68, 69] was used to manually correct any errors in the ER segmentations. The distances of the peptide exit sites of all ribosomes to the closest ER membrane were calculated using a custom python script in 6 out of 9 tomograms that contained ER. Based on a histogram of these distances, a cut-off distance of 8 nm was used to classify ribosomes as being associated with the ER or being cytosolic. We computed the surface area of the ER using the RegionFeaturesJ plugin [70] in ImageJ and used this number to calculate the number of translating ribosomes per ER surface area. The ER-associated ribosome surface densities were computed as *ρ*_ER_ = *N*_ER-assoc_*/A*_ER_, where *N*_ER-assoc_, where NER-assoc is the number of ER-associated ribosomes and AER is the segmented ER surface area. Since the tomograms contained predominantly pmaER, the number of ER-associated ribosomes from the above-described analysis was used to determine the density of pmaER-associated ribosomes. For the cecER and tubER, we adapted the values proportionally based on the surface area estimates described in a previous study on resin embedded samples [24] and the density quantified for the pmaER as described here. Finally, ribosomes were randomly assigned to lattice sites according to the calculated surface areas of the respective ER regions, taking into account that the pmaER membrane proximal to the cell membranes is not decorated with ribosomes.

#### 2.2.6 Proteomics Data Processing and Model Integration

Raw quantitative proteomics data generated by Paulo et al. [37] were obtained from the ProteomeXchange repository (dataset ID PXD002875). These data were generated using a TMT-9plex analysis of *S. cerevisiae* grown on glucose, galactose, and raffinose [37]. Briefly, the authors report that cells were cultured in minimal medium with raffinose, transferred to the respective carbon sources, grown to an OD_600_ of approximately 0.6, lysed by mechanical disruption, and proteins were extracted by chloroform–methanol precipitation, digested with LysC and trypsin, labeled with TMT reagents, fractionated by basic pH reversed-phase chromatography, and analyzed by LC–MS^3^. The raw files were processed using MaxQuant (v2.7.4.0) [71] against the *S. cerevisiae* reference proteome (UniProt UP000002311, 2025-09-29). Absolute protein abundances were estimated using the Proteomic Ruler approach [72], which infers total protein mass from histone signal intensity relative to genome size. Three biological replicates were available for each carbon source; replicate identity was verified using *GAL1* (YBR020W)/*GAL2* (YLR081W) expression for galactose samples and *SUC2* (YIL162W) abundance for raffinose samples, and the mean value per condition was used for downstream analyses. The processed proteomic data were subsequently used to quantify fold changes in *GAL1* and *GAL2* protein abundances when switching from raffinose to galactose, which increased approximately 45-fold and 57-fold, respectively. Processed quantitative results are provided in the Supplementary Information, and analysis scripts are available in Github repository [73].

#### 2.2.7 Accounting for Ribosome Competition in *GAL* Gene Translation

Having determined the total numbers of free cytosolic ribosomes and ER-associated ribosomes, we were able to assign gene transcripts to either ribosome pool based on their predicted site of translation. Generally, we assumed that ribosomes are allocated to mRNAs in proportion to each transcript’s relative abundance within the total cellular mRNA pool. For example, if a given transcript constitutes 5% of the total mRNA population, we assume that 5% of ribosomes are available to translate that transcript. Transcript abundances were obtained by computing the mean of three biological replicates grown in galactose medium [74]. Next, genes annotated with the GO Slim terms “extracellular region” (GO:0005576), “cell wall” (GO:0005618), or “plasma membrane” (GO:0005886) in the Saccharomyces Genome Database [75] were classified as being translated on ER-associated ribosomes; all other genes were assigned to cytosolic ribosomes. Effective cytosolic ribosome numbers were estimated by scaling the total ribosome abundance according to the relative mRNA frequency in each compartment. Using this approach, the combined translation of Gal1p, Gal3p, Gal4p, and Gal80p accounted for 1.2% of cytosolic ribosomes (4,152 ribosomes). To estimate the pool of *effective* ER-associated ribosomes available for co-translational insertion, we first considered proteomic evidence indicating that ER translocon capacity increases upon induction. Proteome-wide measurements showed that Sec61Y abundance increases by approximately 81% when cells are shifted from raffinose to galactose medium [37], suggesting an expansion in the number of ribosomes that can be productively engaged in ER-associated translation. As a baseline, we estimated the total ER-associated ribosome pool—defined here as ribosomes translating ER-targeted transcripts regardless of instantaneous translocon engagement—using transcriptomics of the fraction of ER-translating mRNAs relative to total mRNA abundance [74]. This corresponds to 12.3% of total ribosomes, or approximately 43,788 ribosomes per cell. We then scaled this baseline pool to reflect increased translocon availability under galactose conditions. Finally, within the induced ER-associated translation pool, 12.9% of ribosomes were estimated to translate *GAL2* mRNA based on the relative change in *GAL2* transcript abundance compared with the total change in ER-associated transcripts between raffinose and galactose [74]. Combining these factors yields an estimated 4,256 effective ER-associated ribosomes dedicated to *GAL2* translation per cell, corresponding to approximately 9.7% of the ER-associated ribosome pool and about 1.2% of all cellular ribosomes.

#### 2.2.8 Description of the Extended Yeast Cell Geometry

Our extended yeast cell geometry is shown in Fig 3. This base geometry consisted of 9 spatial regions/compartments (extracellular region, cell wall, plasma membrane, cytosol, nuclear envelope, nucleoplasm, mitochondria, vacuole, and cytosolic ribosomes) and was integrated into the LM software [39] to perform spatially-resolved stochastic simulations with the size of 192×192×192 cubes^3^. For simulations including the ER, the reconstructed geometry included 15 spatial regions (addition of cecER, cecER ribosomes, tubER, tubER ribosomes, pmaER and pmaER ribosomes).

### 2.3 Using the Multi-GPU Solver to Increase RDME Simulation Speed

Compared to prokaryotic systems, modeling yeast cells is substantially more computationally demanding because of their larger size and increased complexity. To address this, we developed a custom multi-GPU solver in Lattice Microbes, `MGPUMpdRdmeSolver`, extending the existing `MpdRdmeSolver` described by Roberts and Hallock et al. [15, 38]. The simulation volume was partitioned along the Z-axis into independent subdomains, each processed by a dedicated GPU. Neighboring GPUs exchanged boundary-layer data through peer-to-peer (P2P) transfers over NVLink or PCIe, minimizing host-memory overhead.

To further improve efficiency, asynchronous CUDA streams were used to overlap computation and communication, while custom packing routines compressed boundary updates into contiguous buffers to reduce data transfer. Affinity mapping allowed us to bind a GPU to a dedicated CPU thread synchronized with `pthread` barriers and `mutexes` to prevent race conditions. Affinity mapping is now introduced and incorporated into Lattice Microbes v2.5.

We benchmarked our implementation on Delta (A100) and Delta AI (H100) supercomputers [76], achieving up to a 2.5-fold speedup using four GPUs (Fig. 4B). This performance enables simulations of larger yeast volumes and longer biological timescales while preserving spatial resolution.

**Figure 4.**
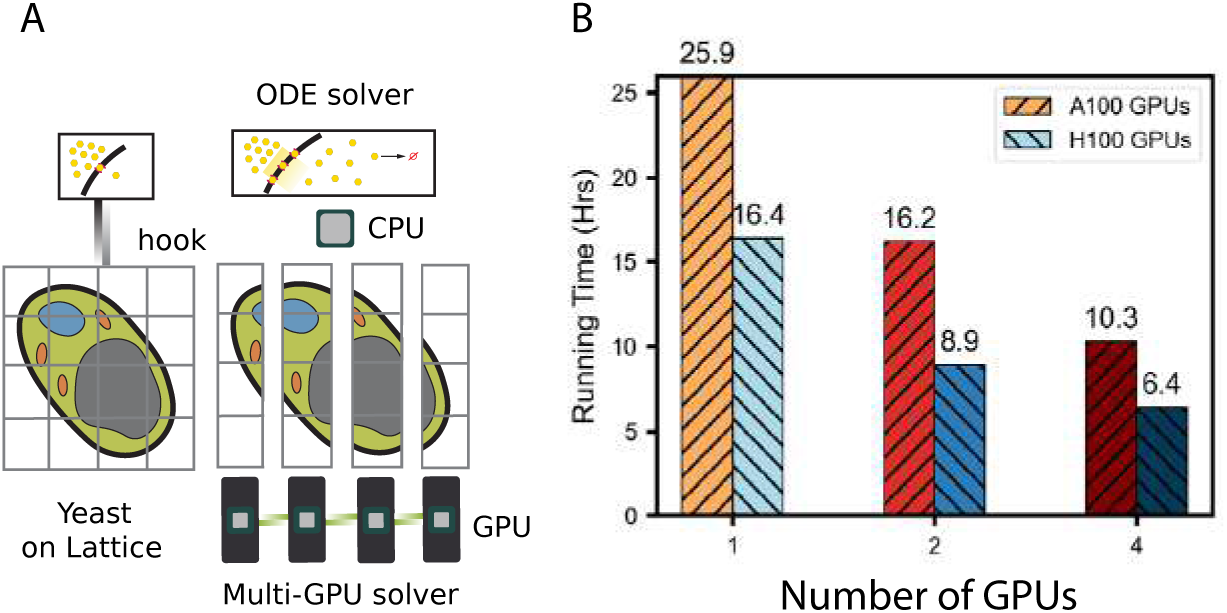
(A) Schematic of multi-GPU domain decomposition used in the yeast RDME simulations. The 3D cellular lattice is partitioned into four subdomains, each assigned to a separate GPU. (B) Performance benchmark per biological hour of 1-, 2-, and 4-GPU simulations on A100 and H100 GPUs, showing sublinear scaling with up to 2.5× speedup on four GPUs.

Each GPU independently processed its assigned subdomain, exchanging only the necessary boundary data during synchronization phases. To optimize runtime performance, we leverage asynchronous CUDA streams that overlap computation with communication in the overlapping regions. Peer-to-peer (P2P) memory transfers via NVLink or PCIe further accelerated data exchange between GPUs by avoiding costly host memory transfers. Moreover, custom packing and unpacking routines compress boundary updates into contiguous buffers, thereby reducing data movement. Fig 4 illustrates how our four-GPU framework partitioned the simulation space and exchanged boundary area data.

### 2.4 Simulation Design and Analysis of Spatially-Resolved Galactose Switch Dynamics

We performed four analyses to isolate how chemical and spatial constraints shape galactose-switch dynamics. First, to test the influence of cell size and diffusion, we compared a previously published CME–ODE model [35] against our base spatial RDME–ODE model, run without chromosome or ER. Owing to stochasticity in both frameworks, we ran 50 replicates for the CME–ODE model and 5 replicates for all RDME–ODE simulations, recording per-replicate counts for all 37 species as well as replicate means and min-max for each model.

Second, to quantify the effect of chromosome localization, we compared the base RDME–ODE model to a genome-resolved RDME–ODE model in which the spatial positions of all 16 yeast chromosomes were explicitly represented (Fig. S2). Third, to assess the influence of ER organization, we compared the genome-resolved (“chromosome”) RDME–ODE model to an extended model that additionally incorporated explicit ER geometry (the “ER model”). In the ER model, *GAL2* mRNA is translation limited to only ER-associated ribosomes. Fourth, we assumed that ribosomes are allocated to mRNAs in proportion to each transcript’s relative abundance and restricted the number of ER-ribosomes available to translate galactose-switch mRNAs accordingly to reflect competition between these transcripts and other cellular mRNAs for a shared ribosome pool.

Across all simulations we recorded abundances for all chemical species, but we emphasized model differences using time courses of *GAL2* -related species (*GAL2*, *GAL2* mRNA, Gal2p, and intracellular galactose). We plotted trajectories for all model iterations and tested for differences at selected time points using Welch’s t-tests; results for all *GAL*-related species are provided in Supplementary File 1 (Sections 3.1–3.4). In all comparisons, kinetic and diffusion parameters were held constant. Simulations were performed under experimentally relevant conditions (11.1 mM extracellular galactose; 0.2% w/v) [18] and were designed to represent a 35.7 fL haploid *S. cerevisiae* cell in G1.

## 3 Results

### 3.1 Cell Geometry and Encounter-Limited Regulation Shift Early Promoter Activity and Accelerate Gal2p-Dependent Uptake

Spatial constraints in eukaryotic cells can alter regulatory outcomes by making low-copy DNA–protein interactions encounter-limited within confined nuclear volumes. To isolate this effect, we first compared the previously published well-stirred CME–ODE model [35] with a baseline spatial RDME–ODE model. This first RDME–ODE iteration incorporated realistic cellular dimensions and cryo-ET–derived major organelles (nucleus, vacuole, mitochondria), thereby enforcing diffusion boundaries and compartmentalized reaction volumes. This baseline spatial model did not yet include explicit chromosome topology, ER geometry, or effective partitioning of the 354,000 ribosomes (Supplementary Video 2:).

Spatial organization and subcellular compartmentalization alone was sufficient to shift early *GAL2* promoter dynamics toward de-repression, producing earlier mRNA accumulation and faster emergence of transporter-linked uptake phenotypes. To quantify these deviations, we used the galactose transporter Gal2p as a representative downstream readout (Fig. 5). Although *GAL2* was initialized in the inhibited state in both models, RDME–ODE simulations exhibited earlier and more sustained activation. Beginning at 7.2 min, *GAL2* showed a higher probability of transitioning to—and remaining in—the activated state in the RDME–ODE framework relative to the CME–ODE framework (*p* ≤ 0.05; Fig. 5A). Consistent with this promoter shift, *GAL2* mRNA levels became significantly elevated after 8.1 min (*p* ≤ 0.05; Fig. 5B). These transcriptional differences propagated to the protein level: plasma membrane–localized Gal2p significantly diverged beginning at 12.3 min (*p* ≤ 0.05; Fig. 5C). As expected for a transporter-limited uptake regime, increased Gal2p delivery was accompanied by enhanced intracellular galactose accumulation, with significance attained after 25.1 min (*p* ≤ 0.01; Fig. 5D).

**Figure 5.**
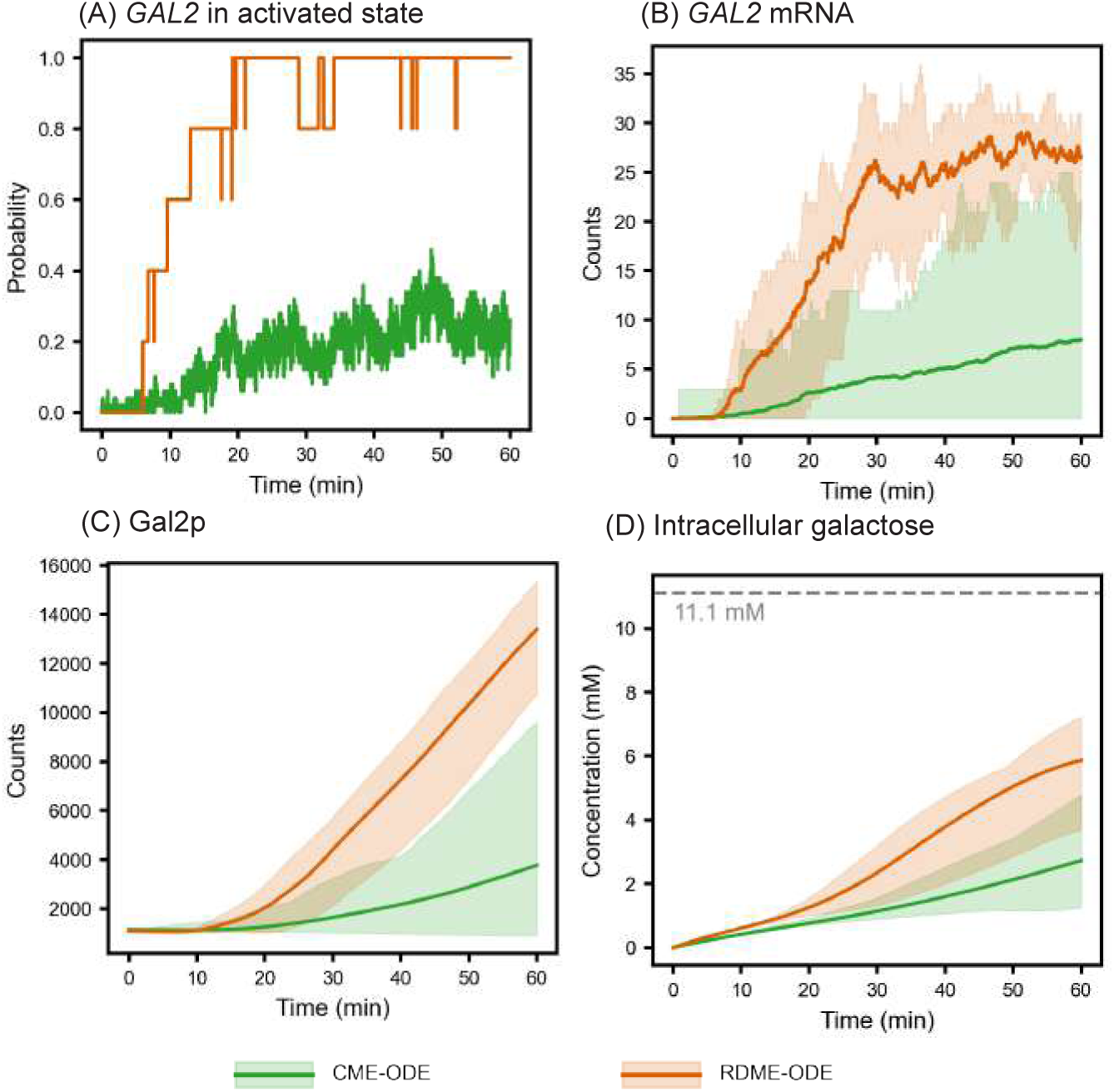
Comparison of *GAL2* -related species between the baseline spatial RDME-ODE model and the CME-ODE model [35]. (A) Probability of *GAL2* gene activity. (B) Abundance of total *GAL2* mRNA. (C) Abundance of Gal2p at the plasma membrane in RDME-ODE simulations, or total Gal2p abundance in CME-ODE simulations. (D) Total intracellular galactose concentration (sum of free and protein-bound intracellular galactose). Solid lines are mean values and shaded areas are the min-max region of 5 replicates (RDME-ODE) or 50 replicates (CME-ODE). The extracellular galactose concentration (11.1 mM) is indicated by the dotted gray line.

Together, these results show that imposing geometry and diffusion constraints can shift induction trajectories on the timescale of minutes to tens of minutes. The differences between the well-stirred CME-ODE models and spatially-resolved RDME-ODE modes arise because when repression involves low-copy repressor acting on single genomic loci(*GAL2*), enforcing spatial locality reduces effective repression events and biases the system toward earlier onset of activation, which then propagates forward to Gal2p accumulation and intracellular galactose uptake (A more detailed explanation is provided in Discussion 4.2). Mechanistically, this baseline divergence from Ramsey’s ODE model [18](Table.2 line CME-ODE and RDME-ODE) motivates to incorporate the following chromosome, ER and ribosome competition.

### 3.2 Inclusion of Chromosome Topology Maintains Final Gal2p Abundance

Chromosome topology could influence nuclear regulation by altering regulator mobility, locus accessibility, or encounter statistics for low-copy repressors. Therefore, we embedded a previously published three-dimensional chromosomal model of *S. cerevisiae* into the nucleoplasm [46] within the RDME–ODE framework. Unlike the baseline RDME–ODE configuration (Sec. 3.1), in which genes were randomly assigned to nucleoplasmic positions, each gene in the chromosome-inclusive model was constrained to its annotated chromosomal start site [75]. The chromosome lattice sites introduced diffusion obstacles within the nucleoplasm, allowing us to directly assess how chromosome topology influences system-level dynamics.

As shown in Fig. 6A and B, both the probability of *GAL2* being in the activated state and the total *GAL2* mRNA abundance were statistically indistinguishable between the chromosome-inclusion and the no-chromosome models (*p >* 0.05 across all time points). Similarly, the abundance of Gal2p in the plasma membrane (Fig. 6C) and the total intracellular galactose concentration (Fig. 6D) did not differ significantly between conditions (*p* = 0.09 at 10 min; *p* = 0.19 at 30 min; *p* = 0.20 at 60 min for Gal2p; *p* = 0.05, *p* = 0.20, and *p* = 0.72 for intracellular galactose, respectively).

**Figure 6.**
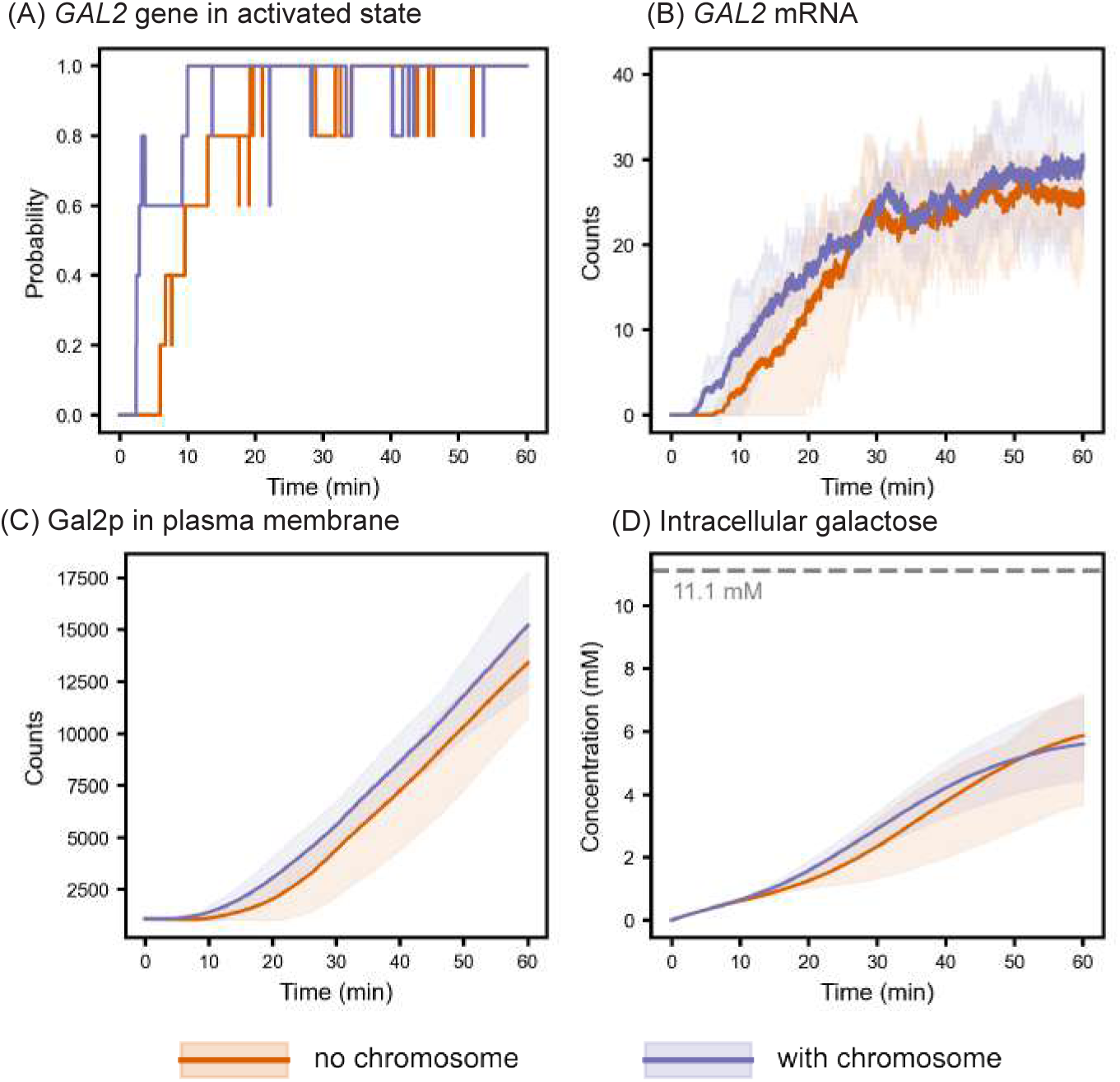
Comparison of *GAL2* -related species between the initial no-chromosome RDME-ODE results and the chromosome-inclusive RDME-ODE results. (A) Probability of *GAL2* being in the activated state. (B) Abundance of total *GAL2* mRNA. (C) Abundance of galactose transporter Gal2p in the plasma membrane. (D) Total intracellular galactose concentration (sum of free and protein-bound intracellular galactose). Solid lines are mean values and shaded areas are the min-max region of 5 replicates (RDME-ODE). The extracellular galactose concentration (11.1 mM) is indicated by the dotted gray line.

Chromosome topology could substantially reduce regulator mobility, altered locus accessibility, or create persistent nuclear gradients that change encounter rates. Here, nucleoplasmic concentrations of both activators and repressors were statistically indistinguishable between conditions (*p >* 0.05; Fig. S10A,B), consistent with chromosome occupancy being too small to measurably perturb effective nuclear exploration at the diffusivities used here. This provides a mechanistic explanation for why chromosome positioning produced little to no change in *GAL2* activation, Gal2p dynamics, or intracellular galactose accumulation in this baseline regime.

### 3.3 Inclusion of ER Geometry Reduces Gal2p Translation Efficiency and Delays Membrane Localization

To increase the spatial complexity of the model, we next incorporated the ER as a distinct cellular compartment. Hereby, we account for the spatial coupling between translation and protein targeting to the plasma membrane [77]. Proteins destined for secretion or membrane insertion are synthesized exclusively by ribosomes associated with the ER, allowing their nascent polypeptide chains to be co-translationally translocated into the ER lumen or membrane via the Sec61 translocon [78]. As an integral plasma membrane protein, Gal2p belongs to this class of proteins substrates and is accordingly synthesized at ER-bound ribosomes [79]. To parametrize ER-associated translation, we determined in Table 1 the ribosome surface densities on ER membranes based on cryo-electron tomograms of S. cerevisiae as described in Methods 2.2.5).

**Table 1.** Comparison of ribosome surface densities on ER regions as calculated in previous literature and in the present study. (Units: ribosomes/*µm*^2^)

| ER Region | Literature Range [24] | Present Study |
| --- | --- | --- |
| cecER | 600–1100 | 424 |
| pmaER | 550–900 | 362 |
| tubER | 250–400 | 162 |

Representative tomographic slices shown in Figure 7 from Supplementary Video 5:) reflect the spatial organization of the ER network within yeast cells into distinct ER subdomains – specifically the central cisternal ER (cecER), the tubular ER (tubER) and the plasma membrane-associated ER (pmaER)(Fig.7A-C). Here, we quantified pmaER associated ribosome densities and proportionally extrapolated them to the cecER and tubER using the relative ribosome association ratios reported in a previous study based on resin embedded samples (Table 1 Literature Range) [24]. Ribosome densities reported there were higher than those measured here by cryo-ET (Table 1 Present Study), likely due to differences in defining ER-associated ribosomes: the present study only counts ribosomes actively translating into the ER based on proximity and orientation, whereas the previous study included all ribosomes within 5 nm regardless of their orientation [24]. Next, we converted ribosome densities were converted into an absolute ER-bound ribosome pool by multiplying the density by the total ER surface area in the geometry used in the simulations (*N_ER−ribo_* = *ρ_ER_/A_ER_*), yielding 7,715 ER-associated ribosomes per cell (Methods 2.2.5). This number is well below the reported median cellular abundance of Sec61 (17,267 copies per cell from SGD), the essential core component of the ER translocation channel [53], indicating that the Sec61 abundance is sufficient to accommodate the observed number of ER-bound ribosomes.

**Figure 7.**
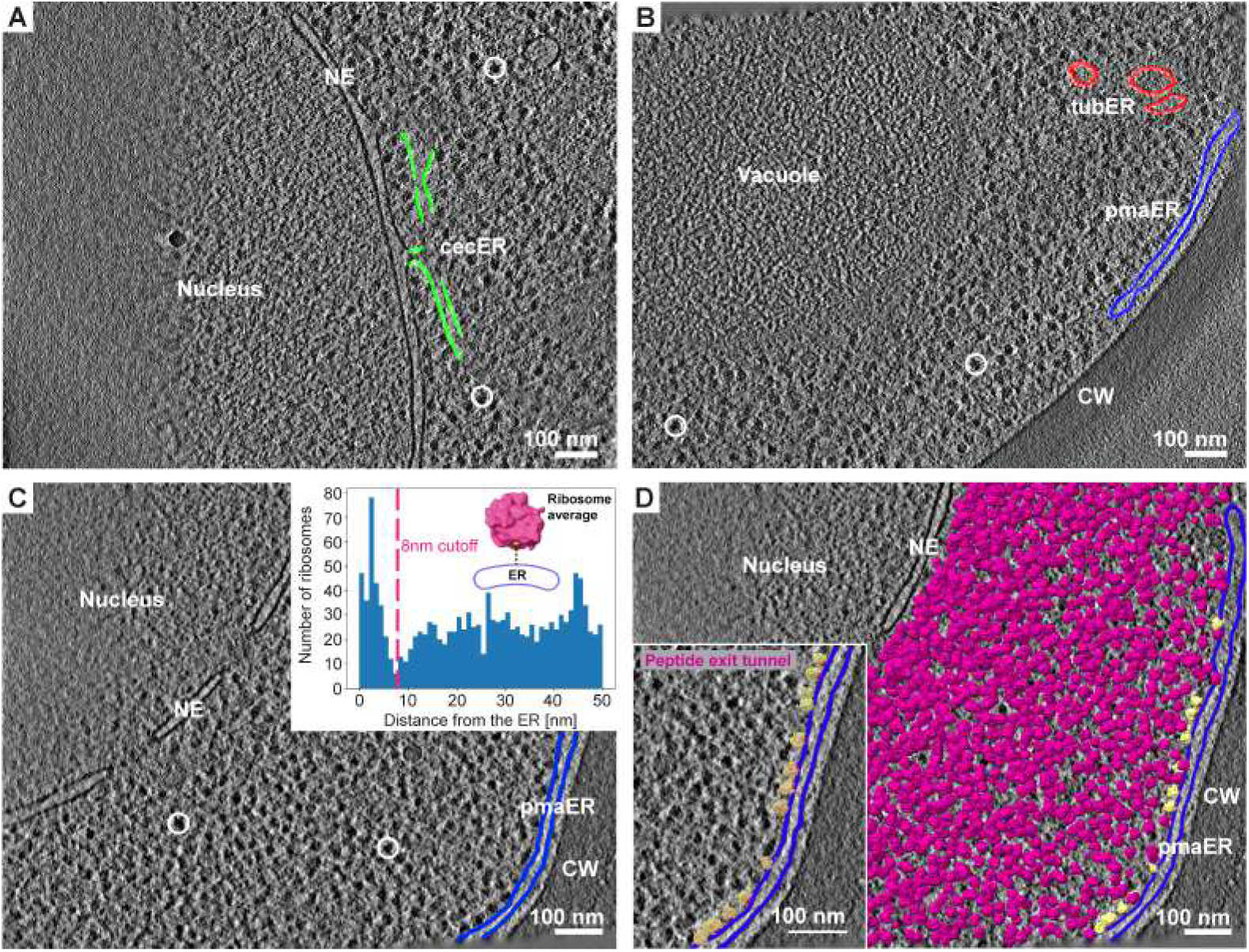
Quantification of ER-associated ribosomes based on cryo-ET data. Representative tomographic slices of *S. cerevisiae* cell depicting the cytosol crowded with ribosomes (dense globular particles), the ER, the cell wall (CW), the vacuole and the nucleus with the nuclear envelope (NE). See movies S 5.1-3 for corresponding tomograms and segmentations. (A) 2D tomographic slice showing cisternal ER (cecER) continuous with NE. Examples of individual ribosomes are encircled in white.

We next asked how this spatial organization affects GAL2 translation by adding ER geometries and restricting GAL2 translation to ER-associated ribosomes. The spatial coupling significantly reduced the fraction of actively translating GAL2 mRNAs, which decreased from 93.12% in the “no-ER” model to 73.48% in the “with ER” model after approximately 15 min (*ρ* ≤ 0.05; Fig. 8A). Inclusion of ER compartments led to a transient accumulation of Gal2p within the ER lumen between 7.4 and 28.7 min (*ρ* ≤ 0.05; Fig. 8B), consistent with delayed ER processing. While early delivery of Gal2p to the plasma membrane was similar in both models, plasma membrane Gal2p levels began to diverge after 11.4 min (*ρ* ≤ 0.05; Fig. 8C). Despite these effects on translational efficiency and subcellular Gal2p distribution, mean intracellular galactose concentrations did not differ significantly between the two models over the simulated timescale (*ρ >* 0.05; Fig. 8C), indicating that ER geometry modulates protein synthesis and trafficking kinetics without substantially altering overall galactose transport flux. Time-resolved spatial trajectories for the “with ER” model are shown in Supplementary Video S5.

**Figure 8.**
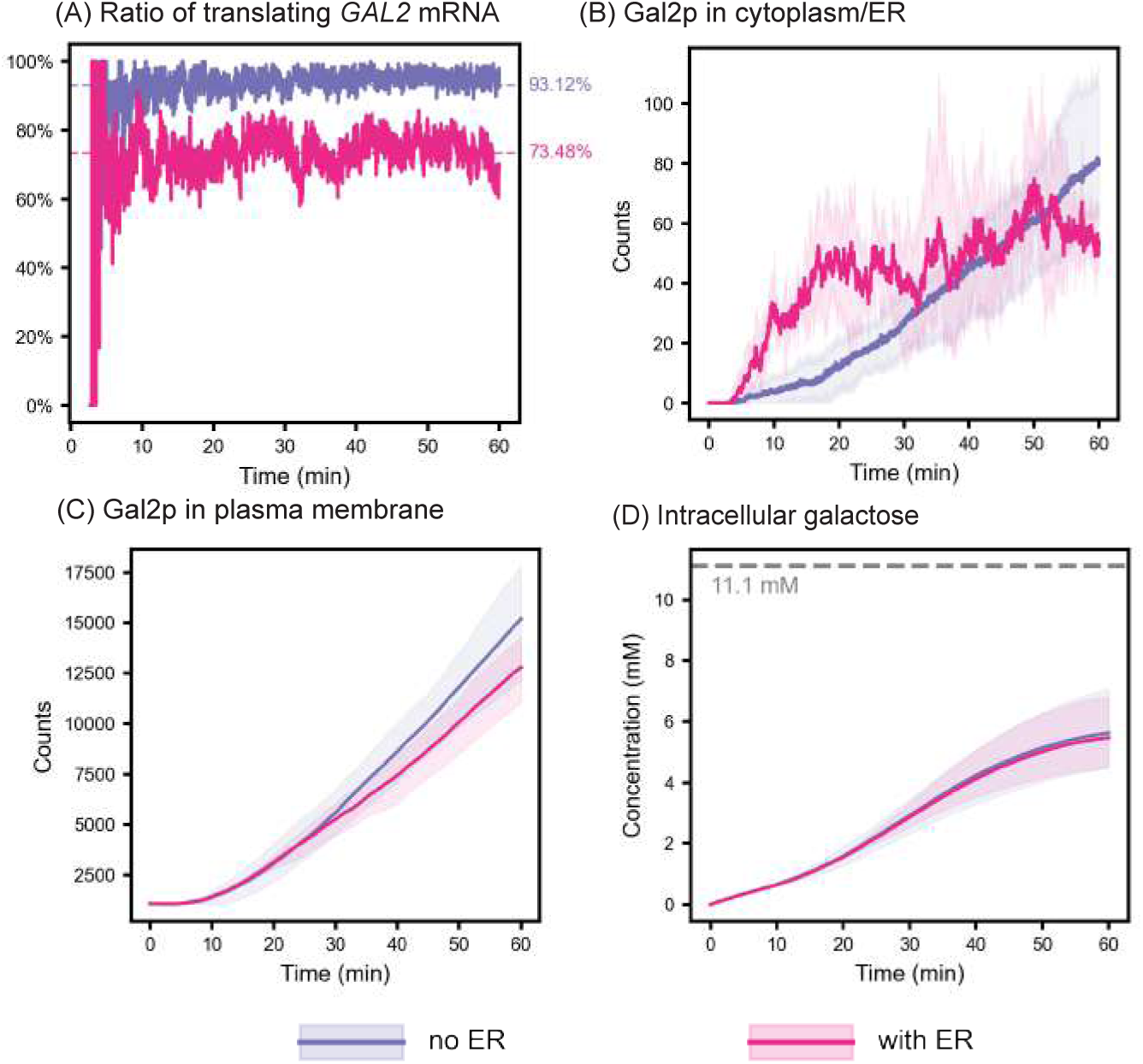
Comparison of *GAL2* –related species between the chromosome-only model (no ER) and ER-inclusive model (with ER). (A) Ratio of ribosome-bound (translating) *GAL2* mRNA to total *GAL2* mRNA. (B) Abundance of Gal2p localized exclusively to the cytoplasm in the no-ER model and to the ER in the ER-inclusive model. (C) Abundance of Gal2p localized to the plasma membrane. (D) Total intracellular galactose concentration (sum of free and protein-bound intracellular galactose). Solid lines are mean values and shaded areas are the min-max region of 5 replicates (RDME-ODE). The extracellular galactose concentration (11.1 mM) is indicated by the dotted gray line.

Tomogram not included in ribosome surface density analysis. See movie S 5.1. (B) 2D tomographic slice containing plasma membrane-associated ER (pmaER) and tubular ER (tubER). Tomogram not included in ribosome surface density analysis. See movie S 5.2. (C) 2D slice of a tomogram included in the ribosome surface density analysis containing pmaER. See movie S5.3. Inset: Subtomogram average of the 80S ribosome with peptide exit site (centroid of the yellow sphere) used to calculate the distance of all ribosomes to the nearest segmented ER membranes and the resulting histogram. A cut-off distance of 8 nm is used to identify translating ribosomes on the surface of the ER. (D) Example of comprehensive 3D annotation of all 80S ribosomes (magenta) in a tomogram. The pmaER is segmented in blue. Ribosomes that have an exit site positioned less than 8 nm from the closest ER membrane are displayed in yellow. Inset: magnification of the pmaER region in (D) showing the orientation of the ribosomes (yellow) peptide exit sites (red spheres) with respect to the ER membrane.

### 3.4 Competition for Ribosomes Creates Translational Bottlenecks that Constrain Gal2p Accumulation in the Plasma Membrane

Cellular induction capacity is constrained by finite translational resources, so promoter activation does not necessarily translate into proportional protein accumulation when mRNAs compete for the free ribosomes. To capture this effect, we restricted the pool of effective ribosomes to account for competition between *GAL*-related transcripts and the broader cellular mRNA pool. These simulations (“ER eff ribo”) were compared with the earlier ER-inclusive model (“ER”). Time-resolved spatial trajectories for the ER-effective ribosome model are provided as Supplementary Video 4:.

Under ribosome-limited conditions, the mean fraction of *GAL2* transcripts bound to ribosomes after 46.3 minutes decreased significantly from 74.48% to 41.68% (*p* ≤ 0.05; Fig. 9A). Spatial analysis of all translating ribosomes (Fig. 9B) showed that, on average, ribosomes were positioned 0.18 *µ*m farther from the nuclear center in the “ER effective-ribo” simulations. Although this analysis includes all translation events rather than *GAL2* -specific ribosomes, the outward shift suggests a general redistribution of translational activity toward the peripheral ER. Reduced ribosome engagement should constrain how efficiently promoter activation is converted into plasma-membrane transporter abundance. Consistent with this expectation, plasma-membrane Gal2p abundance became significantly lower in the “ER eff ribo” case beginning at 12.9 min (*p* ≤ 0.05; Fig. 9C), yielding roughly 50% less Gal2p at the membrane. Intracellular galactose accumulation began to diverge at approximately 25 min (*p* ≤ 0.05; Fig. 9D), with the ribosome-limited case showing reduced accumulation. Together, these results show that translational budget can override transcriptional advantages by limiting Gal2p membrane accumulation and reducing uptake.

**Figure 9.**
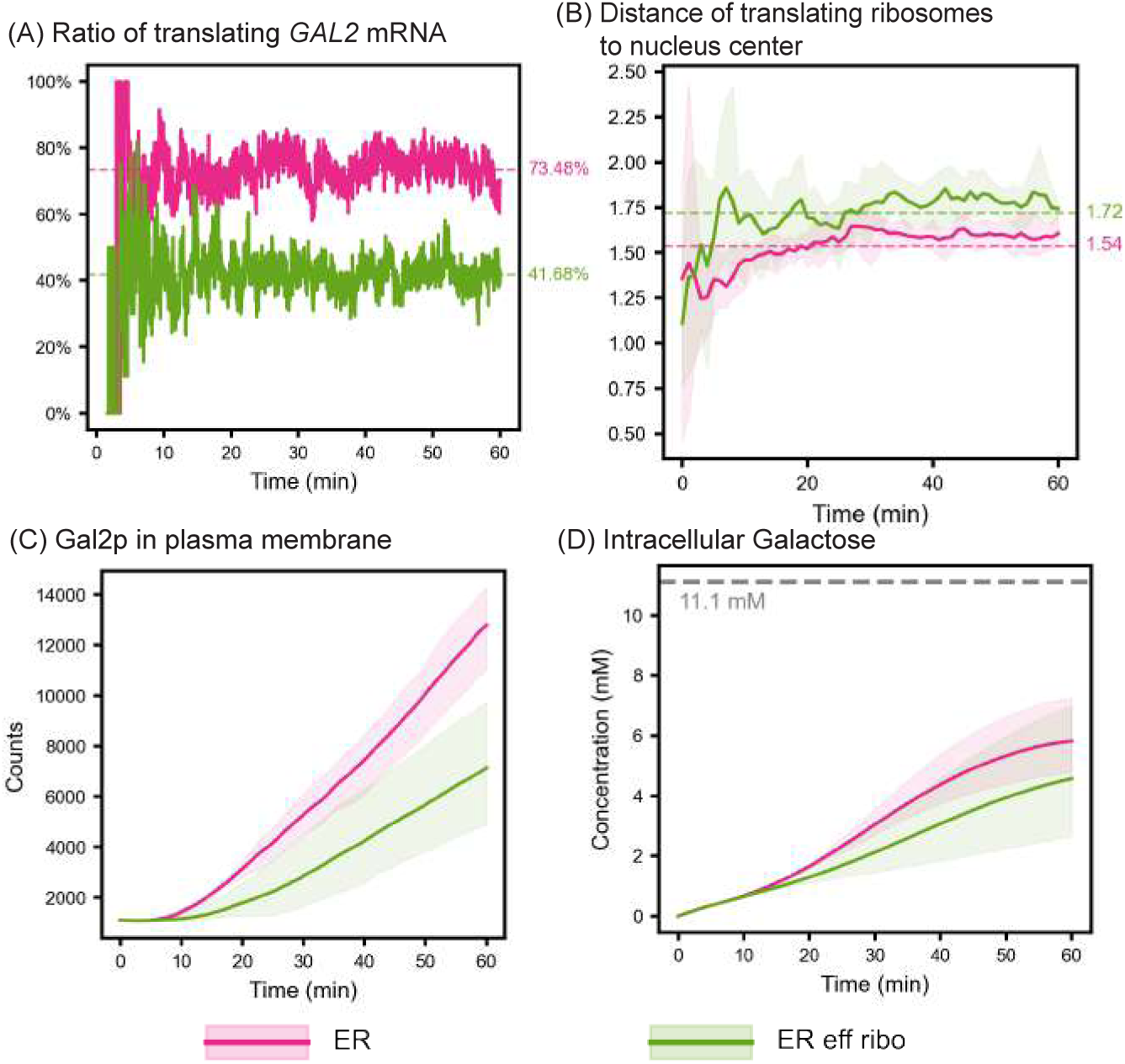
Effects of limiting ribosome availability on *GAL2* -related species. (A) Ratio of actively translating *GAL2* mRNA. (B) Average distance of translating ribosomes to the center of nucleus. (C) Abundance of Gal2p in the plasma membrane. (D) Total intracellular galactose concentration (sum of free and protein-bound intracellular galactose). Solid lines are mean values and shaded areas are the min-max region of 5 replicates (RDME-ODE). The extracellular galactose concentration (11.1 mM) is indicated by the dotted gray line..

### 3.5 Model Validation Against Prior Experiments and Models

Common rate-limiting steps in the galactose switch system are extracellular galactose level. To evaluate whether our modeling frameworks produce realistic quantitative predictions, we compared simulated *GAL* protein induction to two independent published studies that were not used to calibrate our spatial architecture: (i) the time-resolved induction experiments and ODE model of Ramsey et al. [18] and (ii) quantitative proteomics measurements reported by Paulo et al. [37]. Unless otherwise noted, fold changes were computed as the ratio of induced (galactose) to basal (raffinose) protein abundance.

#### 3.5.1 Ribosome-competition models explain most low-galactose induction results reported by Ramsey et al

Ramsey et al. [18] quantified galactose switch induction by transferring *MEL1* -deficient yeast grown in 2% (w/v) raffinose to galactose-containing media ranging from 0.01% to 0.2% (w/v) extracellular galactose concentrations (0.555–11.1 mM). They reported time-course measurements for 0.1% and 0.2% galactose and calibrated their ODE model to 0.1% galactose before evaluating predictions at 0.2%.

To enable direct comparison across frameworks, we evaluated fold changes at 60 minutes under 0.2% (w/v) extracellular galactose (11.1 mM), matching the Ramsey evaluation condition. Table 2 summarizes fold changes in Gal1p, Gal2p, Gal3p, and Gal80p predicted by Ramsey’s ODE model, our well-stirred CME–ODE model, and the successive spatial RDME–ODE variants. Overall, our final spatially informed model (**RDME–ODE(EFF+ER+Chromo)**) produced Gal1p and Gal2p fold changes most similar to Ramsey’s predictions, whereas Gal3p and Gal80p were better approximated by intermediate model variants (Table 2). These results indicate that the impact of explicit spatial structure on predicted protein induction is protein-dependent, consistent with differences in subcellular localization and regulatory kinetics across *GAL* products.

**Table 2.** Fold change in GAL protein abundance at 11.1 mM galactose (0.2% [w/v]) across modeling frameworks. The fold change closest to Ramsey’s ODE result is highlighted in bold for each species. “Chromo” refers to the simulation case “with chromosome” in Sec. 3.2;”ER+Chromo” refers to the simulation case “with ER” in Sec. 3.3; “EFF+ER+Chromo” refers to the simulation case “ER eff ribo” in Sec. 3.4.

| Model | Gal1p | Gal2p | Gal3p | Gal80p |
| --- | --- | --- | --- | --- |
| Ramsey ODE [18] | 4.18 | 6.71 | 3.42 | 1.44 |
| CME–ODE | 1.85 | 3.26 | 2.40 | **2.08** |
| RDME–ODE | 7.78 | 11.59 | 5.55 | 4.12 |
| RDME–ODE(Chromo) | 10.42 | 13.15 | 3.83 | 4.14 |
| RDME–ODE(ER+Chromo) | 7.81 | 11.07 | **3.54** | 3.71 |
| RDME–ODE(EFF+ER+Chromo) | **4.93** | **6.18** | 1.05 | 2.54 |

#### 3.5.2 Model captures most GAL protein abundance changes at high galactose regime

In contrast to Ramsey et al. [18], Paulo et al. [37] reported substantially larger induction fold changes for *GAL* proteins, including approximately 45× (Gal1p) and 57× (Gal2p) when comparing 2% galactose to 2% raffinose, with similarly elevated fold changes for Gal3p and Gal80p. Moreover, baseline protein abundances under raffinose conditions differed markedly between Paulo et al. and the initial protein levels assumed in the Ramsey model, likely reflecting differences in strain background, proteomic quantification methodology, and growth history prior to induction. Because the Ramsey-based parameterization provides a self-consistent kinetic reference framework, we retained those initial conditions for our primary simulations while using the Paulo et al. dataset as an independent benchmark for fold-change validation (Table S17).

Because simulating long time horizons under 2% (w/v) galactose in the stochastic CME/RDME frameworks is computationally prohibitive, we instead constructed a simplified fully deterministic ODE model using the same kinetic parameters as our well-stirred model. We simulated 420 minutes under 2% (w/v) extracellular galactose (111 mM) and compared predicted induction fold changes to both Ramsey’s model and Paulo’s experimental measurements. As shown in Table 3, our deterministic ODE simulation more closely matched the magnitude of Gal1p, Gal2p, and Gal3p induction reported by Paulo et al., whereas Gal80p was substantially overpredicted. This divergence likely reflects parameter-regime mismatch: the regulatory kinetics used in our formulation were calibrated under low galactose conditions (0.2% [w/v]) [35], and applying the same regulation strengths at much higher galactose levels can reduce effective repression of *GAL80*, elevating its predicted steady-state abundance.

**Table 3.** Fold change in total GAL protein abundance at 111 mM galactose (2% [w/v]). Values show the ratio of induced (111 mM galactose) to basal (raffinose) protein levels from Paulo et al. [37], Ramsey et al. [18], and our simplified ODE simulations. Bold values indicate the model result closest to the experimental data.

| Species (Total) | Paulo’s Experiment | Ramsey’s Model | Our Model |
| --- | --- | --- | --- |
| Gal1p | 45 | 11.78 | **55.75** |
| Gal2p | 57 | 16.00 | **45.25** |
| Gal3p | 13 | 6.34 | **15.37** |
| Gal80p | 12 | **2.64** | 30.39 |

Comparisons with Paulo’s experiments indicate our model of regulation is more accurate across different galactose concentrations. But both ours and Ramsey’s original model fail to capture the fold change of Gal80p, suggesting we need more careful and systematic modeling of the galactose switch to make it suitable across scales of external galactose concentrations. Additionally, Palme et al. suggest that there may be a direct inhibitory link between Gal1p and Gal80p, which could be included in future versions of the model [80].

## 4 Discussion

A central goal of computational cell biology is to develop mechanistic models capable of predicting cellular phenotypes and responses across diverse experimental conditions. In eukaryotic systems, this requires accounting not only for biochemical regulation but also for intracellular spatial organization, which constrains diffusion, local concentrations, and access to subcellular machinery. Leveraging decades of biochemical characterization together with imaging-derived cellular geometries, we demonstrate that explicitly representing spatial heterogeneity can quantitatively alter regulatory dynamics: even a relatively simple module—the yeast galactose switch—exhibits substantially different behavior when modeled within a spatially organized intracellular context.

We systematically increased the spatial heterogeneity and complexity in our model to determine the effect each addition had on the cell responses to galactose uptake. First, we converted a well-stirred model of the yeast galactose switch network [35] into a spatially informed framework using a discretized lattice representation of the cell. Voxels were treated as locally well mixed, reactions occurred only among co-localized molecules, and diffusion followed predefined rules between neighboring sites. Next, we incorporated experimentally informed geometrical representations of subcellular organelles into our model and constrained integral membrane proteins (i.e., Gal2p) to be translated only on ER-associated ribosomes. Finally, we accounted for competition between *GAL*-related transcripts and all other mRNAs by using a limited ribosome pool. These spatial considerations stand in stark contrast to traditional well-stirred computational models, which allow all molecules to react regardless of their spatial positions. By enforcing locality (i.e., requiring molecules to be in physical proximity to interact) and grounding the intracellular architecture in experimental measurements, our framework moves closer to a physically realistic environment in which to study regulatory dynamics. The following subsections aim to describe and add context to each of these progressive steps.

### 4.1 Comparison of Well-Stirred and Spatially Informed Simulations - Figures 5 and S3

Our spatially informed models exhibited a decrease in the overall rate of second-order elementary reactions relative to well-stirred models (Figure 5A and Figure S3). This effect arises from two interrelated consequences of spatial locality in RDME simulations: (i) bimolecular reactions require co-localization, so reactants must first diffuse into the same lattice site before reacting; and (ii) reaction propensities are computed from voxel-local copy numbers rather than global abundances, such that typical bimolecular propensities (∝ *n_A_*(**x**)*n_B_*(**x**)) are reduced when molecules are distributed across many voxels (check Supplementary File 1: section S5 for detailed derivation). Additionally, explicit compartmentalization alters molecular fluxes and imposes compartment-specific initial molecular abundances, further modulating reaction statistics and regulatory dynamics.

#### 4.1.1 Differences in Compartment-Specific Initial Molecular Abundances

The *initial* spatial distribution of molecules imposed by compartment volumes was an additional difference between spatially informed and well-stirred simulations. Because initial molecular counts were allocated according to cytosolic and nucleoplasmic volume fractions, the nucleoplasmic abundance of G80d in our spatially informed models was lower at *t* = 0 than in the well-stirred model, which assumed uniform mixing across the entire intracellular volume. This reduced initial nuclear repressor availability increased the probability of early promoter activation, contributing to the enhanced gene activation observed in spatially informed simulations.

#### 4.1.2 Search Times Decrease the Possibility of Second-Order Reaction Events

The inclusion of spatial heterogeneity introduces the need for reacting molecules to find one another. Because our models include only elementary reactions, first-order reactions (single-reactant) are not affected by this requirement, whereas second-order reactions depend on molecular encounter times. These encounter limitations are most pronounced when one reactant has low copy number, such as gene regulation by the repressor G80d, where each target locus exists in a single copy.

To demonstrate these effects, consider the repression reaction:

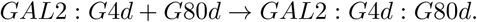

To estimate the mean time for a single G80d molecule to encounter the *GAL2* locus, we model its nuclear exploration as an ergodic random walk on a finite lattice. The average time a repressor needs to leave and return to the GAL2 gene site is proportional to the number of search sites (*N*_nuc_) divided by the number of target sites (*N_GAL2_*). If the motion of the repressor is diffusive, then it has a time step of 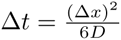 and the overall time to revisit the *GAL2* locus is:

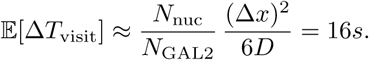

Although this value is small, it is in sharp contrast to the implicit instantaneous encounter assumption of well-stirred models. As a result, spatial search times reduce the effective frequency of repression events, thereby contributing to the increased probability of promoter activation in spatially informed simulations.

#### 4.1.3 Enforcing Molecular Locality Decreases the Effective Rate of Second-Order Reaction Events

In stochastic chemical kinetics, the *bimolecular propensity* refers to the probability per unit time that a second-order reaction will occur, typically computed from the copy numbers of two reactant species and a reaction-rate constant. In a well-stirred CME–ODE framework, bimolecular propensities are calculated from global copy numbers under the assumption of instantaneous mixing within the nuclear volume. In contrast, RDME simulations impose well-mixed conditions only within individual voxels: bimolecular reactions can occur only when reactants occupy the same voxel, and the corresponding propensity depends on local copy numbers. Consequently, spatial discretization reduces the *effective* system-level frequency of second-order reaction events by (i) requiring co-localization and (ii) replacing global molecule counts with typically smaller local counts.

This effect can be illustrated for the repression reaction

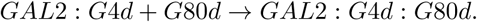

In a CME model, the propensity scales with the global nuclear abundance of G80d and the cell volume. For example, if the nucleus contains *N_G_*_80*d*_ = 100 copies of G80d and a single copy of *GAL*2 : *G*4*d*, then the well-stirred bimolecular propensity is proportional to *N_G_*_80*d*_. In the RDME framework, however, only the local number of repressors in the voxel containing the *GAL2* locus contributes to the propensity, i.e., *a*_RDME_ ∝ *N_G_*_80*d*:vox_, where *N_G_*_80*d*:vox_ denotes the number of G80d molecules in the same voxel as *GAL*2 : *G*4*d*. Because molecules are distributed across many voxels, *N_G_*_80*d*:vox_ ≪ *N_G_*_80*d*_ on average, yielding lower local propensities and fewer repression events over time.

The same encounter limitations apply to activators; however, activator unbinding is ∼10× slower in our model, leading to long promoter residence times once bound. Because the mean bound time scales as *τ*_bound_≈ 1*/k*_off_, a 10-fold decrease in *k*_off_ increases activator residence time by ∼10-fold. As a result, activation is less sensitive to search-time limitations than repression. This interpretation is further supported by a global parameter sensitivity analysis, which identifies nuclear transport/diffusion parameters for G80d as the dominant drivers of Gal2p abundance at 60 min (seeSupplementary File 1: section S5 and Fig. S14).

#### 4.1.4 Asymmetries in Molecular Fluxes Into and Out of the Nucleus Result in Decreased Gene Activation Rates

In addition to reducing second-order reaction events, introducing spatial structure also changed the effective exchange of regulators across the nuclear envelope. In the well-stirred CME–ODE model, nuclear import and export are represented with symmetric, high transport rates, implicitly assuming rapid equilibration between nucleus and cytosol. In spatial RDME simulations, however, transport becomes a *search problem*: a molecule must diffuse to a nuclear pore to cross the envelope. This immediately introduces an asymmetry, because escaping from the nucleus requires searching only the nuclear volume, whereas entering from the cytosol requires searching a much larger volume. We therefore interpret the altered nuclear fluxes using classical first-passage arguments [81, 82].

For a diffusing species such as G80d, the mean time to *exit* the nucleus through nuclear pores can be approximated as

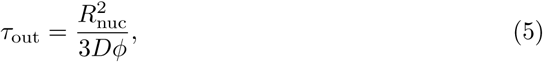

where *R*_nuc_ is the nuclear radius, *D* is the diffusion coefficient, and *ϕ* is the fraction of the nuclear surface occupied by pores. Using our parameters and *ϕ* ≈ 0.06, this corresponds to an effective export rate of *k*_out_ ≈ 11.03 min*^−^*^1^.

In contrast, the mean time for *entry* from the cytosol is dominated by the larger cytoplasmic search volume. A simple scaling approximation gives

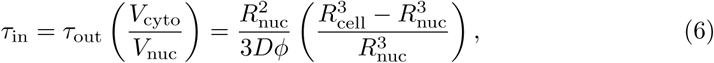

where *R*_cell_ is the cell radius. For a typical yeast geometry (*R*_cell_ ≈ 2*R*_nuc_), *V*_cyto_*/V*_nuc_ ≈ 7, yielding a substantially smaller import rate of *k*_in_ ≈ 1.57 min*^−^*^1^. These physically derived, geometry-dependent rates contrast sharply with the symmetric transport rates assumed in the well-stirred model (*k*_in_ = *k*_out_ = 500 min*^−^*^1^), highlighting how cellular geometry and localized pores can reshape effective nucleocytoplasmic transport kinetics.

#### 4.1.5 The Effect of Spatial Heterogeneities is Dependent on Reaction Network Topology

The combination of processes described in *Sections 4.1.1-4.1.4* resulted in increased gene activation of all *GAL* genes (other than *GAL4* which was constitutively expressed) when spatial heterogeneities were included in our model (see Figure 5A and Figure S3). Although an increase in gene activation may appear to be a general result of adding spatial heterogeneities to computational models, we argue that this result is specific to the reaction network topology of our current study. One can just as easily imagine a scenario in which a gene becomes activated by association with an activating transcription factor. As this activation reaction would also be second-order, this altered reaction network would produce a decrease in gene activation in the presence of spatial heterogeneities. Therefore, the inclusion of spatial heterogeneities will have varying effects on gene expression based on differences in reaction network topologies.

### 4.2 Effects of Subcellular Organelle Architectures

In addition to the creation of voxels and discrete chemical environments, we also tested the effects of including subcellular organelle architectures on our computational models. In our current model implementations, we investigated the effects of chromosome and ER geometries as well as constraints on ribosome accessibility. Each of these considerations is described below in more detail either focusing on their effects on G80d/gene activation or on Gal2p production.

#### 4.2.1 Inclusion of Chromosomal Geometries Did Not Significantly Alter Gene Activation - Figure 6

Incorporating chromosome geometries could, in principle, alter the time for G80d to locate *GAL* loci by changing the accessible nucleoplasmic volume, repositioning genes, or introducing local excluded volume near loci. However, we observed no statistically significant changes in *GAL* gene activation upon adding chromosome geometry (Fig. S4), indicating that any such effects are negligible under the present parameter regime.

A simple diffusivity estimate provides a quantitative rationale. In our model, chromosome lattice sites occupy only ∼3.2% of the nuclear volume. For diffusion in a spherical domain containing randomly distributed impermeable obstacles with volume fraction *ϕ* ≈ 0.032, percolation-inspired scalings [83] predict only a modest reduction in the effective diffusion coefficient,

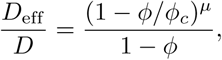

which yields *D*_eff_ */D* ≈ 0.97 using parameters for approximately spherical obstacles (*ϕ_c_* = 0.672, *µ* = 1.35) [84]; treating obstacles as more cylinder-like gives a similar estimate (*D*_eff_ */D* ≈ 0.95). Such small changes in nucleoplasmic diffusivity are consistent with the absence of measurable differences in activation dynamics.

Consistent with this interpretation, varying gene placement also had little effect in our simulations. When chromosome sites were removed and genes were placed either centrally or randomly throughout the nucleus, gene activation remained statistically indistinguishable between scenarios (Fig. S13). This insensitivity likely reflects the relatively fast diffusion of G80d in the nucleoplasm: when search is rapid compared to the timescale of promoter binding/unbinding, geometric repositioning has limited leverage. We therefore expect gene positioning to become more influential in regimes with slower nucleoplasmic diffusion or stronger crowding.

Finally, local excluded volume near loci was also insufficient to measurably perturb activation. Even genes with relatively high nearby chromosome occupancy (e.g., *GAL1* and *GAL2*, ∼23.1% of nearest-neighbor sites occupied) did not differ significantly from genes with near-zero local occupancy (e.g., *GAL3* and *GAL80*; Table S13), and the same held when considering two- and three-step neighborhoods (Table S15). In the present model, chromosome positions are static and local occupancy fractions therefore remain constant over time, further limiting potential effects.

More substantial chromosome-driven effects may emerge when additional sources of nuclear crowding are included, such as chromatin-associated proteins (e.g., histones and chromatin-binding factors) that occupy a significant fraction of nuclear volume but are not explicitly represented here.

#### 4.2.2 ER-Associated Ribosome Requirements for Membrane Proteins Decreased the Production of Gal2p - Figure 8

Introducing the ER into the simulations coupled two distinct constraints on membrane-protein synthesis: (i) *capacity* —only ER-associated ribosomes were permitted to translate Gal2p, reducing the effective ribosome pool available for Gal2p production; and (ii) *localization*—*GAL2* mRNA had to diffuse to ER-associated sites and be translated there, rather than on randomly distributed cytosolic ribosomes. When both constraints were enforced simultaneously, Gal2p abundance decreased, but the initial simulations did not resolve the relative contributions of reduced ribosome number versus enforced ribosome location.

To disentangle these effects, we performed targeted ablation simulations that independently varied ribosome *number* and ribosome *placement*. First, to isolate ribosome *number* effects without introducing ER geometry, we reduced the cytosolic ribosome count (345,964) for *GAL2* mRNA to match the ER-associated ribosome count (7,833) while leaving ribosomes randomly distributed in the cytosol. This reduction produced a substantial drop in Gal2p output (Fig. S14A), indicating that limited translational capacity alone can strongly suppress Gal2p production. Next, holding the reduced ribosome count fixed as 7,833, we compared random placement to ER-like placement by positioning the same 7,833 ribosomes at the ER-associated locations used in the ER-constrained model. Under ER-like placement, Gal2p output increased relative to the random-placement scenario (Fig. S14B), implying that ribosome spatial clustering partially compensates for reduced ribosome number. Finally, Comparison the diffusion of Gal2p via the geometry of ER lumen v.s. the 3D cytosol, where the diffusion coefficents are kept the same, showed Gal2p production was not significantly altered, suggesting that the dominant effects arise from ribosome availability and spatial clustering rather than diffusion via the geometry of ER lumen.

Together, these ablations indicate that the ER effect on Gal2p is not explained solely by ribosome count or solely by spatial localization, but by their combined influence: restricting translation to a smaller ribosome pool decreases Gal2p, whereas concentrating that pool into ER-localized sites increases local translation efficiency, consistent with a clustering-driven “hotspot” effect. This behavior is compatible with ER-associated polysome-like organization, where ribosome proximity can increase the likelihood of repeated translation of nearby transcripts and promote bursty, sustained protein production.

In contrast, other *GAL* gene products showed no significant changes upon enforcing ER-ribosome requirements because they are not integral membrane proteins and were therefore not subject to ER-localized translation constraints. Their dynamics were influenced only by the modest change in the cytosolic ribosome pool (from 345,964 to 338,131; Tables S2 and S3), which produced at most minor effects.

#### 4.2.3 Ribosomal Competition Decreased Protein Levels of All *GAL* Genes - Figure 9

Guided by transcriptomic comparisons (see Methods 2.2.7), restricting the pool of ribosomes available to translate *GAL*-related mRNAs reduced protein output across all *GAL* genes. This trend is consistent with the targeted ablation simulations described in the above section, Fig. 9 and Supplementary File 1: 3.4, which showed that limiting translational capacity alone is sufficient to lower steady-state protein abundance. Importantly, these differences could not be explained by changes in mRNA levels (Fig. S5), indicating that the observed effects arise primarily from altered translation efficiency rather than transcriptional regulation. Together, these results highlight how spatial and resource constraints on ribosomes can quantitatively reshape gene-expression outcomes beyond transcriptional control.

### 4.3 Limitations and Future Directions

The yeast model presented here integrates experimentally derived spatial architectures with reaction–diffusion dynamics and deterministic kinetics, but several limitations remain. First, spatial simulations are computationally demanding, restricting both simulated biological timescales and the number of stochastic replicates, which may limit the detection of rare or highly variable molecular trajectories. Second, our spatial architectures are based on static structural snapshots and therefore do not capture dynamic organelle remodeling or vesicle-mediated trafficking [85]. For example, Gal2p delivery to the plasma membrane was approximated through ER diffusion without explicitly modeling SRP-dependent targeting or vesicular transport, which may influence translation localization and protein abundance. Third, only a reduced representation of the Leloir pathway was included, potentially affecting predicted intracellular galactose levels and downstream regulatory responses. Finally, several post-transcriptional and post-translational processes that shape mRNA and protein abundances are represented only implicitly, limiting quantitative accuracy of absolute molecular copy numbers. Future spatial whole-cell models that incorporate dynamic organelle organization, trafficking pathways, and expanded metabolic and regulatory networks will further improve predictive fidelity.

### 4.4 Conclusions

Overall, we have demonstrated that spatial heterogeneity critically alters predictions made by computational eukaryotic cell models. By coupling structural, geometrical, and molecular features into a single spatially-resolved model, we show how seemingly modest changes in diffusion, encounter probabilities, and molecular accessibility can accumulate into substantial differences in gene activation, transporter production, and nutrient uptake. Our work provides an important first step toward the construction of a spatially-resolved whole-cell model of budding yeast. In addition, our work highlights both the opportunities and the challenges that arise when extending whole-cell modeling to eukaryotic cells.

## Data and Code Availability

The simulation and analysis code used in this study is available at the GitHub repository [73]. Instructions to reproduce all results are provided in the repository README file. A permanent, citable version of this repository has also been archived on Zenodo (DOI: https://doi.org/10.5281/zenodo.18425352 and https://doi.org/10.5281/zenodo.18433248).

## Acknowledgments

The authors thank David Bianchi and Troy Brier for their advice in using Lattice Microbes. This work used Delta and Delta AI at National Center for Supercomputing Applications (NCSA) through allocation BIO250002 from the Advanced Cyberinfrastructure Coordination Ecosystem: Services & Support (ACCESS) program, which is supported by U.S. National Science Foundation grants #2138259, #2138286, #2138307, #2137603, and #2138296. We also thank our research programmer Henry Li for his performance test on the code on Delta and Delta AI at NCSA. We are grateful to Sara Kathrin Goetz for acquiring the cryo-electron tomograms, the EMBL cryo-EM platform for instrument access, and Thomas Hoffmann and the EMBL IT team for computational support. J.M. was supported by a European Research Council Starting Grant (3DCellPhase 760067), Chan Zuckerberg Initiative grant for visual proteomics (2023-331612) and EMBL. This project was supported by the National Science Foundation Science and Technology Center for Quantitative Cell Biology (DBI-2243257). T.W., M.C.S., and Z.L.S. were also partially supported by NSF DBI-2243257. Research reported in this publication by Z.R.T. was supported by the Cancer Center at Illinois–Beckman Institute Postdoctoral Fellows Program, sponsored by the Cancer Center at Illinois and the Beckman Institute for Advanced Science and Technology, University of Illinois Urbana–Champaign. The content is solely the responsibility of the authors and does not necessarily represent the official views of the program sponsors.

## 5 Supporting Information

### Supplementary File 1

**Simulation parameters and supplementary results.** This file provides detailed information on species names, reactions, and simulation parameters (including kinetic rates and diffusion coefficients). It also contains additional simulation results referenced in the main text.

### Supplementary Video 1

**Final ER Geometry.** This video shows the final ER geometry used in the present model.

### Supplementary Video 2

**RDME–ODE simulation trajectory (60 min, normal geometry).** This video shows the 60-minute trajectory of the RDME–ODE simulation under the normal geometry condition (no chromosomes, ER or effective ribosomes).

### Supplementary Video 3

**RDME–ODE simulation trajectory (60 min, ER-included geometry).** This video shows the 60-minute trajectory of the RDME–ODE simulation under the ER-inclusive geometry condition.

### Supplementary Video 4

**RDME–ODE simulation trajectory (60 min, ER geometry with effective ribosomes).** This video shows the 60-minute trajectory of the RDME–ODE simulation under the ER-inclusive geometry condition with effective ribosome modeling.

### Supplementary Video 5

**Cryo-ET movies of wild-type *S. cerevisiae*.**

*Video S5.1:* Cryo-ET of wildtype *S. cerevisiae* showing the nucleus with the nuclear envelope, and the central cisternal ER (cecER) that is continuous with the NE. The cytosol is crowded with electron dense ribosomes. Scale bar: 100 nm.

*Video S5.2:* Cryo-ET of wildtype *S. cerevisiae* showing the cytosol crowded with electron dense ribosomes, the plasma membrane-associated ER (pmaER) in blue and the tubular ER (tubER) in red. Scale bar: 100 nm.

*Video S5.3:* Cryo-ET of wildtype *S. cerevisiae* with 3D annotations of peripheral ER (pmaER, blue) and cytosolic ribosomes (magenta). Ribosomes with peptide exit tunnels closer than 8 nm to the ER are colored in yellow with the peptide exit tunnels indicated by magenta spheres. Scale bar: 100 nm.

## Notes

https://github.com/Luthey-Schulten-Lab/Yeast_Galactose_Switch_RDMEODE

https://doi.org/10.5281/zenodo.18425352

https://doi.org/10.5281/zenodo.18433248

### Competing Interest Statement

The authors have declared no competing interest.

### Summary of Updates

Final update before submission to IPOLS computational biology

