## Supplemental File 1 for "Spatially Resolved Reaction–Diffusion Modeling Reveals Effects of Intracellular Spatial Heterogeneity on Yeast Galactose Network Dynamics"

### Supplementary Information for Spatial Heterogeneity Alters the Dynamics of the Yeast Galactose Switch: Insights from 4D RDME–ODE Hybrid Simulations

Tianyu Wu[1,2], Marie-Christin Spindler[3,2], Abner T. Apsley[5], Emmy Earnest[4], Zane R. Thornburg[5,6], Julia Mahamid[3,7,2*], Zaida Luthey-Schulten[8,2,1*]

**1** Center for Biophysics and Quantitative Biology, University of Illinois at Urbana–Champaign, Urbana, IL, USA
**2** National Science Foundation Science and Technology Center for Quantitative Cell Biology, Beckman Institute for Advanced Science and Technology, University of Illinois at Urbana–Champaign, Urbana, IL, USA
**3** Molecular Systems Biology Unit, European Molecular Biology Laboratory (EMBL), Heidelberg, Germany
**4** Department of Physics, University of Illinois at Urbana–Champaign, Urbana, IL, USA
**5** Beckman Institute for Advanced Science and Technology, University of Illinois at Urbana-Champaign, Urbana, IL, USA
**6** Cancer Center at Illinois, University of Illinois at Urbana-Champaign, Urbana, IL, USA
**7** Cell Biology and Biophysics Unit, European Molecular Biology Laboratory (EMBL), Heidelberg, Germany
**8** Department of Chemistry, University of Illinois at Urbana–Champaign, Urbana, IL, USA

* *

3.9 Ribosome Number and Spatial Organization Jointly Limit GAL2 Translation 23

**4 Comparison of Predicted and Measured Final Fold Changes of GAL Proteins Across Experimental and Modeling Studies 23**

**5 Comparison of propensity of second order reaction in CME and RDME 24**

**6 Mathematical interpretation of ER-dependent translation and trafficking 25**

**7 Sensitivity Analysis on kinetic parameters 26**

### 1 Species, reactions and parameters in the simulation

#### 1.1 Species

**Table S1.** Species and their annotations

| Species | Annotation |
|---|---|
| $DG_1$ | gene encoding Gal1 with nothing bound (free) |
| $DG_1 : G_{4d}$ | gene encoding Gal1 bound to Gal4 dimer(activated) |
| $DG_1 : G_{4d} : G_{80d}$ | gene encoding Gal1 bound to the Gal4 dimer and Gal80 dimer(repressed) |
| $DG_2$ | gene encoding Gal2 with nothing bound(free) |
| $DG_2 : G_{4d}$ | gene encoding Gal2 bound to the Gal4 dimer(activated) |
| $DG_2 : G_{4d} : G_{80d}$ | gene encoding Gal2 bound to the Gal4 dimer and Gal80 dimer(repressed) |
| $DG_3$ | gene encoding Gal3 with nothing bound (free) |
| $DG_3 : G_{4d}$ | gene encoding Gal3 bound to the Gal4 dimer(activated) |
| $DG_3 : G_{4d} : G_{80d}$ | gene encoding Gal3 bound to the Gal4 dimer and Gal80 dimer(repressed) |
| $DG_4$ | gene encoding Gal4 with nothing bound(activated) |
| $DG_{80}$ | gene encoding Gal80 with nothing bound(free) |
| $DG_{80} : G_{4d}$ | gene encoding Gal80 bound to the Gal4 dimer(activated) |
| $DG_{80} : G_{4d} : G_{80d}$ | gene encoding Gal80 bound to the Gal4 dimer and Gal80 dimer (repressed) |
| $DG_{rep}$ | gene encoding the reporter fluorescent protein with nothing bound(free) |
| $DG_{rep} : G_{4d}$ | gene encoding reporter bound to the Gal4 dimer(activated) |
| $DG_{rep} : G_{4d} : G_{80d}$ | gene encoding reporter bound to the Gal4 dimer and Gal80 dimer(repressed) |
| $R_1$ | mRNA for Gal1 |
| $R_2$ | mRNA for Gal2 |
| $R_3$ | mRNA for Gal3 |
| $R_4$ | mRNA for Gal4 |
| $R_{80}$ | mRNA for Gal80 |
| $R_{rep}$ | mRNA for the reporter gene |
| $G_1$ | galactokinase metabolizes galactose |
| $G_2$ | galactose transporter |
| $G_3$ | galactose sensing transcription factor |
| $G_{3i}$ | Gal3 bound to a galactose molecule |
| $G_{4d}$ | the transcriptional activator |
| $G_4$ | the monomer of the transcriptional activator |
| $G_{80d}$ | the transcriptional repressor dimer |
| $G_{80}$ | the monomer of the transcriptional repressor |
| $G_{80d} : G_{3i}$ | Gal80 dimer bound to Gal3i; the transcriptional repressor sequestered in the cytoplasm |
| $G_{rep}$ | a yellow/green fluorescence reporter protein (YFP) |
| GAE | extracellular galactose |
| GAI | intracellular galactose |
| $G_2GAI$ | galactose bound to the Gal2 transporter on the intracellular side |
| $G_2GAE$ | galactose bound to the Gal2 transporter on the extracellular side |
| $G_1GAI$ | galactose bound to the Gal1 transporter on the intracellular side |

#### 1.2 Initial counts of species

| Species | State | Region | Count |
|---|---|---|---|
| DG1 | DG1_G4d_G80d | nucleoplasm | 1 |
| DG2 | DG2_G4d_G80d | nucleoplasm | 1 |
| DG3 | DG3_G4d_G80d | nucleoplasm | 1 |
| DG80 | DG80_G4d_G80d | nucleoplasm | 1 |
| DGrep | DGrep_G4d_G80d | nucleoplasm | 1 |
| DG4 | DG4 | nucleoplasm | 1 |
| R1 | | cytoplasm | 0 |
| R2 | | cytoplasm | 0 |
| R3 | | cytoplasm | 1 |
| R4 | | cytoplasm | 0 |
| R80 | | cytoplasm | 1 |
| Rrep | | cytoplasm | 0 |
| G1 | | cytoplasm | 132 |
| G2 | | Plasma membrane | 1,156 |
| G3 | | cytoplasm | 4,341 |
| G4d | | nucleoplasm | 308 |
| Grep | | cytoplasm | 132 |
| G80 | | cytoplasm | 0 |
| G80 | | nucleoplasm | 0 |
| G80d | | cytoplasm | 157 |
| G80d | | nucleoplasm | 157 |
| ribosome | | ribosome | 345,964 |

**Table S2.** Species initial counts in RDME for no-chromosome and with-chromosome cases. If not mentioned, the initial counts are 0. Species counts are rounded to integers from Ramsey *et al.* [1].

| Species | State | Region | Count |
|---|---|---|---|
| ribosome | | ribosome | 0 |
| cytosolic ribosome | | ribosome | 338,131 |
| ER-associated ribosome | | ribosome | 7,833 |

**Table S3.** Change in species initial counts when switching from the no-chromosome/with-chromosome models to the ER-inclusive case; species not shown remain unchanged.

| Species | State | Region | Count |
|---|---|---|---|
| cytosolic ribosome | | ribosome | 4,152 |
| dummy cytosolic ribosome | | ribosome | 333,979 |
| ER-associated ribosome | | ribosome | 4,256 |
| dummy ER-associated ribosome | | ribosome | 3,577 |

**Table S4.** Changes in initial species counts when transitioning from the ER-inclusive model to the effective-ribosome model. Species not shown remain unchanged. "Dummy" denotes ribosomes that are unavailable for translating galactose-switch mRNAs due to competition from non-switch transcripts.

| Species | Concentration(mol/L) |
|---|---|
| G1 | $7.23 \times 10^{-9}$ |
| G2 | $7.97 \times 10^{-8}$ |
| GAI | 0 |
| G1GAI | 0 |
| G2GAE | 0 |
| G2GAI | 0 |

**Table S5.** Species initial concentration in ODE.

*To be consistent, G1 and G2 concentration is the initial counts in RDME divided by $NA \cdot V$.The species counts are rounded to integers from Ramsey et al. [1].*

#### 1.3 Chemical reactions in the RDME system

These are all reactions included in the RDME system, they are grouped based on their functions (regulation, transcription, translation, mRNA degradation and protein/complex degradation).
*Kinetic parameters are adopted from previously published studies [2].*

**Table S6.** Summary of gene regulation.

| Chemical Reaction | Order | Region | Kinetic Parameters | Comments |
|---|---|---|---|---|
| $D_{G1} + G_{4d} \longrightarrow D_{G1}{:}G_{4d}$ | 2 | nucleoplasm | $3.583 \times 10^{7} \mathrm{M}^{-1}\mathrm{s}^{-1}$ | GAL4 dimer binds to GAl1 gene |
| $D_{G1}{:}G_{4d} \longrightarrow D_{G1} + G_{4d}$ | 1 | nucleoplasm | $6.410 \times 10^{-5}\mathrm{s}^{-1}$ | GAL4 dimer dissociates from Gal1 gene |
| $D_{G1}{:}G_{4d} + G_{80d} \longrightarrow D_{G1}{:}G_{4d}{:}G_{80d}$ | 2 | nucleoplasm | $3.583 \times 10^{5} \mathrm{M}^{-1}\mathrm{s}^{-1}$ | GAL80 dimer binds to GAl1 gene |
| $D_{G1}{:}G_{4d}{:}G_{80d} \longrightarrow D_{G1}{:}G_{4d} + G_{80d}$ | 1 | nucleoplasm | $1.422 \times 10^{-3}\mathrm{s}^{-1}$ | GAL80 dimer dissociates from Gal1 gene |
| $D_{G2} + G_{4d} \longrightarrow D_{G2}{:}G_{4d}$ | 2 | nucleoplasm | $3.583 \times 10^{7} \mathrm{M}^{-1}\mathrm{s}^{-1}$ | GAL4 dimer binds to GAl2 gene |
| $D_{G2}{:}G_{4d} \longrightarrow D_{G2} + G_{4d}$ | 1 | nucleoplasm | $1.684 \times 10^{-3}\mathrm{s}^{-1}$ | GAL4 dimer dissociates from Gal2 gene |
| $D_{G2}{:}G_{4d} + G_{80d} \longrightarrow D_{G2}{:}G_{4d}{:}G_{80d}$ | 2 | nucleoplasm | $3.583 \times 10^{5} \mathrm{M}^{-1}\mathrm{s}^{-1}$ | GAL80 dimer binds to GAl2 gene |
| $D_{G2}{:}G_{4d}{:}G_{80d} \longrightarrow D_{G2}{:}G_{4d} + G_{80d}$ | 1 | nucleoplasm | $2.250 \times 10^{-3}\mathrm{s}^{-1}$ | GAL80 dimer dissociates from Gal2 gene |
| $D_{G3} + G_{4d} \longrightarrow D_{G3}{:}G_{4d}$ | 2 | nucleoplasm | $3.583 \times 10^{7} \mathrm{M}^{-1}\mathrm{s}^{-1}$ | GAL4 dimer binds to GAl3 gene |
| $D_{G3}{:}G_{4d} \longrightarrow D_{G3} + G_{4d}$ | 1 | nucleoplasm | $6.720 \times 10^{-4}\mathrm{s}^{-1}$ | GAL4 dimer dissociates from Gal3 gene |
| $D_{G3}{:}G_{4d} + G_{80d} \longrightarrow D_{G3}{:}G_{4d}{:}G_{80d}$ | 2 | nucleoplasm | $3.583 \times 10^{5} \mathrm{M}^{-1}\mathrm{s}^{-1}$ | GAL80 dimer binds to GAl3 gene |
| $D_{G3}{:}G_{4d}{:}G_{80d} \longrightarrow D_{G3}{:}G_{4d} + G_{80d}$ | 1 | nucleoplasm | $8.842 \times 10^{-3}\mathrm{s}^{-1}$ | GAL80 dimer dissociates from Gal3 gene |
| $D_{G80} + G_{4d} \longrightarrow D_{G80}{:}G_{4d}$ | 2 | nucleoplasm | $3.583 \times 10^{7} \mathrm{M}^{-1}\mathrm{s}^{-1}$ | GAL4 dimer binds to GAl80 gene |
| $D_{G80}{:}G_{4d} \longrightarrow D_{G80} + G_{4d}$ | 1 | nucleoplasm | $6.720 \times 10^{-5}\mathrm{s}^{-1}$ | GAL4 dimer dissociates from Gal80 gene |

| Chemical Reaction | Order | Region | Kinetic Parameters | Comments |
|---|---|---|---|---|
| $D_{G80}{:}G_{4d} + G_{80d} \longrightarrow D_{G80}{:}G_{4d}{:}G_{80d}$ | 2 | nucleoplasm | $3.583 \times 10^5 \mathrm{M}^{-1}\mathrm{s}^{-1}$ | GAL80 dimer binds to GAl80 gene |
| $D_{G80}{:}G_{4d}{:}G_{80d} \longrightarrow D_{G80}{:}G_{4d} + G_{80d}$ | 1 | nucleoplasm | $8.842 \times 10^{-3}\mathrm{s}^{-1}$ | GAL80 dimer dissociates from Gal80 gene |
| $D_{G_{rep}} + G_{4d} \longrightarrow D_{G_{rep}}{:}G_{4d}$ | 2 | nucleoplasm | $3.583 \times 10^7 \mathrm{M}^{-1}\mathrm{s}^{-1}$ | GAL4 dimer binds to GFP/YFP gene |
| $D_{G_{rep}}{:}G_{4d} \longrightarrow D_{G_{rep}} + G_{4d}$ | 1 | nucleoplasm | $6.410 \times 10^{-5}\mathrm{s}^{-1}$ | GAL4 dimer dissociates from GFP/YFP gene |
| $D_{G_{rep}}{:}G_{4d} + G_{80d} \longrightarrow D_{G_{rep}}{:}G_{4d}{:}G_{80d}$ | 2 | nucleoplasm | $3.583 \times 10^5 \mathrm{M}^{-1}\mathrm{s}^{-1}$ | GAL80 dimer binds to GFP/YFP gene |
| $D_{G_{rep}}{:}G_{4d}{:}G_{80d} \longrightarrow D_{G_{rep}}{:}G_{4d} + G_{80d}$ | 1 | nucleoplasm | $1.422 \times 10^{-3}\mathrm{s}^{-1}$ | GAL80 dimer dissociates from GFP/YFP gene |

| Chemical Reaction | Order | Region | Kinetic Parameters | Comments |
|---|---|---|---|---|
| $D_{G_1}{:}G_{4d} \longrightarrow D_{G_1}{:}G_{4d} + R_1$ | 1 | nucleoplasm | $0.012\mathrm{s}^{-1}$ | GAL1 transcription |
| $D_{G_2}{:}G_{4d} \longrightarrow D_{G_2}{:}G_{4d} + R_2$ | 1 | nucleoplasm | $0.042\mathrm{s}^{-1}$ | GAL2 transcription |
| $D_{G_3}{:}G_{4d} \longrightarrow D_{G_3}{:}G_{4d} + R_3$ | 1 | nucleoplasm | $7.110 \times 10^{-3}\mathrm{s}^{-1}$ | GAL3 transcription |
| $D_{G_{rep}}{:}G_{4d} \longrightarrow D_{G_{rep}}{:}G_{4d} + R_{rep}$ | 1 | nucleoplasm | $0.019\mathrm{s}^{-1}$ | YFP/GFP transcription |
| $D_{G_{80}}{:}G_{4d} \longrightarrow D_{G_{80}}{:}G_{4d} + R_{80}$ | 1 | nucleoplasm | $0.010\mathrm{s}^{-1}$ | GAL80 transcription |
| $D_{G_4} \longrightarrow D_{G_4} + R_4$ | 1 | nucleoplasm | $1.650 \times 10^{-4}\mathrm{s}^{-1}$ | GAL4 transcription(no regulation) |

**Table S7.** Summary of transcription reactions in the system. Kinetic parameters are from the previous study [2]

.

**Table S8.** Summary of mRNA degradation

| Chemical Reaction | Order | Region | Kinetic Parameters | Comments |
|---|---|---|---|---|
| $R_1 \longrightarrow \varnothing$ | 1 | cytoplasm; nucleoplasm; ribosomes | $3.727 \times 10^{-4}\mathrm{s}^{-1}$ | R1 degradation |
| $R_2 \longrightarrow \varnothing$ | 1 | cytoplasm; nucleoplasm; ribosomes | $1.284 \times 10^{-3}\mathrm{s}^{-1}$ | R2 degradation |

| Chemical Reaction | Order | Region | Kinetic Parameters | Comments |
|---|---|---|---|---|
| $R_3 \longrightarrow \varnothing$ | 1 | cytoplasm; nucleoplasm; ribosomes | $4.443 \times 10^{-4}\mathrm{s}^{-1}$ | R3 degradation |
| $R_4 \longrightarrow \varnothing$ | 1 | cytoplasm; nucleoplasm; ribosomes | $4.127 \times 10^{-4}\mathrm{s}^{-1}$ | R4 degradation |
| $R_{80} \longrightarrow \varnothing$ | 1 | cytoplasm; nucleoplasm; ribosomes | $4.813 \times 10^{-4}\mathrm{s}^{-1}$ | R80 degradation |
| $R_{rep} \longrightarrow \varnothing$ | 1 | cytoplasm; nucleoplasm; ribosomes | $5.777 \times 10^{-4}\mathrm{s}^{-1}$ | GFP/YFP mRNA degradation |

**Table S9.** Summary of translation.

| Chemical Reaction | Order | Region | Kinetic Parameters | Comments |
|---|---|---|---|---|
| Ribosome + $R_1 \longrightarrow$Ribosome:$R_1$ | 2 | Ribosomes | $3.454 \times 10^5\mathrm{M}^{-1}\mathrm{s}^{-1}$ | R1 binds to Ribosome |
| Ribosome:$R_1 \longrightarrow$Ribosome+ $R_1 + G_1$ | 1 | Ribosomes | $0.032s^{-1}$ | R1 translation |
| Ribosome + $R_2 \longrightarrow$Ribosome:$R_2$ | 2 | Ribosomes | $3.454 \times 10^5\mathrm{M}^{-1}\mathrm{s}^{-1}$ | R2 binds to Ribosome |
| Ribosome:$R_2 \longrightarrow$Ribosome+ $R_2 + G_2$ | 1 | Ribosomes | $0.225s^{-1}$ | R2 translation |
| Ribosome + $R_3 \longrightarrow$Ribosome:$R_3$ | 2 | Ribosomes | $3.454 \times 10^5\mathrm{M}^{-1}\mathrm{s}^{-1}$ | R3 binds to Ribosome |
| Ribosome:$R_3 \longrightarrow$Ribosome+ $R_3 + G_3$ | 1 | Ribosomes | $0.924s^{-1}$ | R3 translation |
| Ribosome + $R_4 \longrightarrow$Ribosome:$R_4$ | 2 | Ribosomes | $3.454 \times 10^5\mathrm{M}^{-1}\mathrm{s}^{-1}$ | R4 binds to Ribosome |
| Ribosome:$R_4 \longrightarrow$Ribosome+ $R_4 + G_4$ | 1 | Ribosomes | $0.178s^{-1}$ | R4 translation |
| Ribosome + $R_{rep} \longrightarrow$Ribosome:$R_{rep}$ | 2 | Ribosomes | $3.454 \times 10^5\mathrm{M}^{-1}\mathrm{s}^{-1}$ | reporter mRNA binds to Ribosome |
| Ribosome:$R_{rep} \longrightarrow$Ribosome+ $R_{rep} + G_{rep}$ | 1 | Ribosomes | $0.096s^{-1}$ | reporter mRNA translation |
| Ribosome + $R_{80} \longrightarrow$Ribosome:$R_{80}$ | 2 | Ribosomes | $3.454 \times 10^5\mathrm{M}^{-1}\mathrm{s}^{-1}$ | R80 binds to Ribosome |
| Ribosome:$R_{80} \longrightarrow$Ribosome+ $R_{80} + G_{80}$ | 1 | Ribosomes | $0.061s^{-1}$ | R80 translation |

**Table S10.** Summary of complex formation

| Chemical Reaction | Order | Region | Kinetic Parameters | Comments |
|---|---|---|---|---|
| $G_4 + G_4 \longrightarrow G_{4d}$ | 2 | cytoplasm; nucleoplasm | $3.583 \times 10^{10}\mathrm{M}^{-1}\mathrm{s}^{-1}$ | GAL4 dimer formation |
| $G_{4d} \longrightarrow G_4 + G_4$ | 1 | cytoplasm; nucleoplasm | $1.667 \times 10^{-5}\mathrm{s}^{-1}$ | GAL4 dimer dissociation |
| $G_{80} + G_{80} \longrightarrow G_{80d}$ | 2 | cytoplasm; nucleoplasm | $3.583 \times 10^{10}\mathrm{M}^{-1}\mathrm{s}^{-1}$ | GAL80 dimer formation |
| $G_{80d} \longrightarrow G_{80} + G_{80}$ | 1 | cytoplasm; nucleoplasm | $1.667 \times 10^{-5}\mathrm{s}^{-1}$ | GAL80 dimer dissociation |
| $G_3 \longrightarrow G_{3i}$ | 1 | cytoplasm | $266.940 \cdot [GAI]\mathrm{s}^{-1}$ | GAL3 binds to GAI |
| $G_{3i} \longrightarrow G_3$ | 1 | cytoplasm | $14.833\mathrm{s}^{-1}$ | GAL3:GAI dissociation |
| $G_{3i} + G_{80d} \longrightarrow G_{80d}{:}G_{3i}$ | 2 | cytoplasm | $9.214 \times 10^{6}\mathrm{M}^{-1}\mathrm{s}^{-1}$ | GAL3:GAI binds to Gal80 dimer |
| $G_{80d}{:}G_{3i} \longrightarrow G_{3i} + G_{80d}$ | 1 | cytoplasm | $2.660 \times 10^{-4}\mathrm{s}^{-1}$ | GAL3:GAI:G80d dissociation |

*Kinetic parameters are adopted from previously published studies [1].*

**Table S11.** Summary of protein/complex degradation

| Chemical Reaction | Order | Region | Kinetic Parameters | Comments |
|---|---|---|---|---|
| $G_{3i} \longrightarrow \varnothing$ | 1 | cytoplasm | $1.925 \times 10^{-4}\mathrm{s}^{-1}$ | GAL3 degradation |
| $G_{80D}{:}G_{3i} \longrightarrow \varnothing$ | 1 | cytoplasm | $9.625 \times 10^{-5}\mathrm{s}^{-1}$ | Gal3:GAI:Gal80 degradation |
| $G_1 \longrightarrow \varnothing$ | 1 | cytoplasm | $6.418 \times 10^{-5}\mathrm{s}^{-1}$ | G1 degradation |
| $G_2 \longrightarrow \varnothing$ | 1 | cytoplasm plasmaMembrane | $6.418 \times 10^{-5}\mathrm{s}^{-1}$ | Gal2 degradation |
| $G_3 \longrightarrow \varnothing$ | 1 | cytoplasm | $1.925 \times 10^{-4}\mathrm{s}^{-1}$ | Gal3 degradation |
| $G_4 \longrightarrow \varnothing$ | 1 | cytoplasm; nucleoplasm | $1.155 \times 10^{-4}\mathrm{s}^{-1}$ | Gal4 degradation |
| $G_{rep} \longrightarrow \varnothing$ | 1 | cytoplasm | $1.925 \times 10^{-4}\mathrm{s}^{-1}$ | Reporter degradation |
| $G_{80} \longrightarrow \varnothing$ | 1 | cytoplasm; nucleoplasm | $1.155 \times 10^{-4}\mathrm{s}^{-1}$ | Gal80 degradation |

#### 1.4 Chemical reactions in the system

**Table S12.** Summary of galactose transportation and metabolism in ODE

| Chemical Reaction | Order | Kinetic Parameters | Comments |
|---|---|---|---|
| $G_1 + GAI \longrightarrow G_1GAI$ | 2 | $1.442 \times 10^5 \mathrm{M}^{-1}\mathrm{s}^{-1}$ | G1 bind to galactokinase |
| $G_1GAI \longrightarrow G_1 + GAI$ | 1 | $30.708\mathrm{s}^{-1}$ | |
| $G_1GAI \longrightarrow G_1$ | 1 | $55.833\mathrm{s}^{-1}$ | metabolite of Galactose |
| $G_2 \longrightarrow G2GAE$ | 1 | $1.123 \times 10^5 \cdot [GAE]\mathrm{s}^{-1}$ | transporter bind to external galactose |
| $G_2GAE \longrightarrow G_2$ | 1 | $39.875\mathrm{s}^{-1}$ | |
| $G_2GAE \longrightarrow G_2GAI$ | 1 | $72.5\mathrm{s}^{-1}$ | symmetric transport |
| $G_2GAI \longrightarrow G_2GAE$ | 1 | $72.5\mathrm{s}^{-1}$ | symmetric transport |
| $G_2GAI \longrightarrow G_2 + GAI$ | 1 | $39.875\mathrm{s}^{-1}$ | release galactose to cytoplasm |

*Kinetic parameters are adopted from previously published studies [2].*

##### 1.5 Diffusion Coefficients for species in RDME and ODE

The diffusion coefficients for all species in the RDME simulation. If the diffusion rule is not mentioned, then it is not allowed between regions.

**Table S13.** Diffusion Coefficients for Various Species between different Regions.

| **Species** | **From** | **To** | **Diffusion Coefficient** $(m^2/s)$ |
|---|---|---|---|
| genes | Any | Any | 0 |
| mRNAs | nucleoplasm | nucleoplasm | $0.05 \times 10^{-12}$ |
| | nucleoplasm | cytoplasm | $0.05 \times 10^{-12}$ |
| | cytoplasm | cytoplasm | $0.05 \times 10^{-12}$ |
| | cytoplasm | ribosome | $0.05 \times 10^{-12}$ |
| | ribosome | cytoplasm | $0.05 \times 10^{-12}$ |
| proteins and complexes | ribosome | cytoplasm | $2.74 \times 10^{-12}$ |
| | cytoplasm | cytoplasm | $1.00 \times 10^{-12}$ |
| | ribosome | ER | $2.74 \times 10^{-12}$ |
| G4/G4d/G80/G80d | nucleoplasm | nucleoplasm | $1.00 \times 10^{-12}$ |
| | nucleoplasm | cytoplasm | $1.00 \times 10^{-12}$ |
| | cytoplasm | nucleoplasm | $1.00 \times 10^{-12}$ |
| G2 | cytoplasm | plasma membrane | $1.00 \times 10^{-12}$ |
| | plasma membrane | plasma membrane | $0.01 \times 10^{-12}$ |
| | ER | ER | $1 \times 10^{-12}$ |
| | ER | cytoplasm | $1 \times 10^{-12}$ |

*Diffusion Coefficients are adopted from previously published studies [3, 4].*

#### 2 Geometry Reconstruction

We demonstrate the process of reconstructing of nuclear envelope as an example:

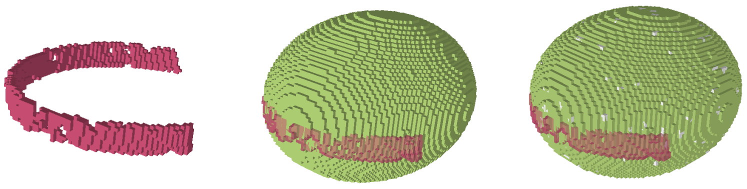

(A) (B) (C)

**Figure S1.** Geometric Reconstruction of the Nuclear Envelope.(A) Segmentation of the nuclear envelope membranes included in cryo-ET data of the yeast cell slice.(B) Ellipsoidal fitting of the nuclear envelope.(C) Addition of nuclear pore complexes as white crosses to the reconstructed envelope.

#### 2.1 Geometric Reconstructions Parameters

Here, we assumed plasma membrane, nucleus envelope and cell wall are ellipsoidal structures and used python *scipy* to fit the parameters. All fitted parameters are listed below:

**Table S14.** Ellipsoid parameters: center coordinates, Euler orientations, and semi-axis lengths. Length unit: lattice coordinates; Angle unit: radians

| **Structure** | **Center** $(x, y, z)$ | **Orientation** $(\alpha, \beta, \gamma)$ | **Semi-axes** $(a, b, c)$ |
|---|---|---|---|
| Plasma Membrane | (96, 96, 96) | (3.39, −1.59, −0.90) | (61, 92, 61) |
| Nuclear Envelope | (132, 81, 112) | (0.16, −1.74, 0.56) | (30, 37, 30) |
| Cell Wall | (96, 96, 96) | (3.39, −1.59, −0.90) | (65, 96, 65) |

Because the mitochondria and vacuole are randomly positioned within the cytoplasmic region with fixed volumetric occupancy, each execution of the script generates distinct mitochondrial and vacuolar configurations with varying parameters. Consequently, these values are not reported in the table.

#### 2.2 Chromosome model

We utilized the static chromosome model [5], applying spatial transformations and uniform scaling to maximize nucleoplasmic occupancy while preventing overlap with the nuclear envelope.

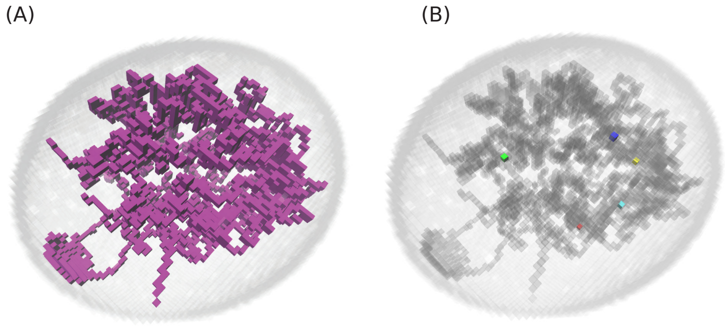

**Figure S2.** (A)Chromosomes in the nuclear envelope. (B) Gene locations in the chromosome. GAL1: cyan, GAL2: red, GAL3: yellow, GAL4: green, GAL80: blue.

| Gene | 1-step | 2-step | 3-step |
|---|---|---|---|
| GAL1 | 23.1% | 18.4% | 8.3% |
| GAL2 | 23.1% | 11.2% | 6.9% |
| GAL3 | 0.0% | 3.1% | 10.6% |
| GAL4 | 0.0% | 8.2% | 8.3% |
| GAL80 | 0.0% | 14.3% | 10.1% |

**Table S15.** Neighbor occupancy by chromosome lattice.

#### 2.3 Nuclear Pore

We measured the surface density of nuclear pore complex to be $13.21/\mu m^2$ based on the cryo-ET data [3]. Then, we assumed the nuclear pores are uniformly distributed on the entire nuclear envelope which make the total number of 139.

### 3 Additional Simulation Results

#### 3.1 Additional *GAL*-related species for Section 3.1

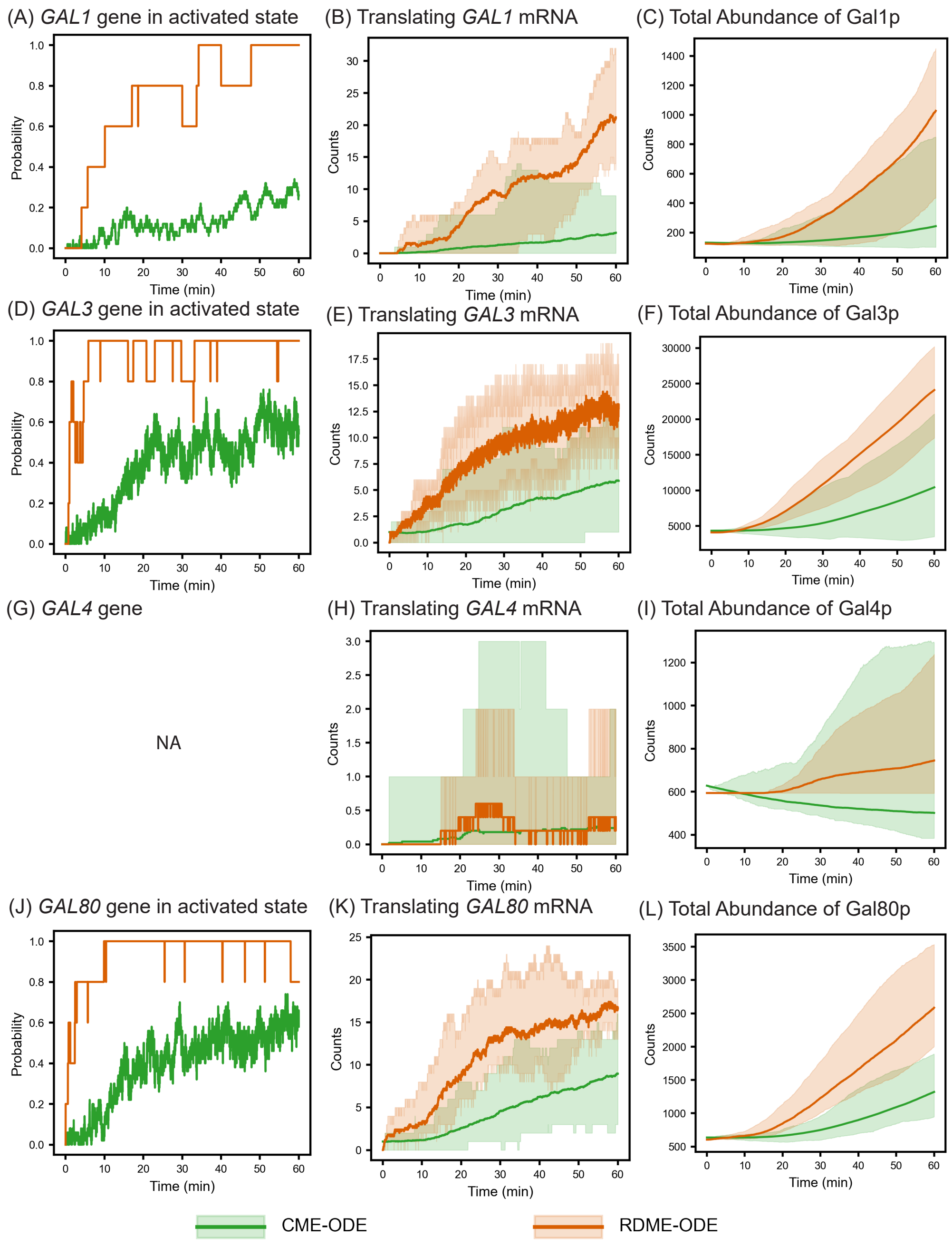

**Figure S3**

The Gal4 gene trajectory (Figure S3G) is not shown because Gal4 transcription is defined as $\varnothing \longrightarrow$ mRNA, meaning that no explicit *GAL4* gene species exists in the CME model.

#### 3.2 Additional *GAL*-related species for Section 3.2

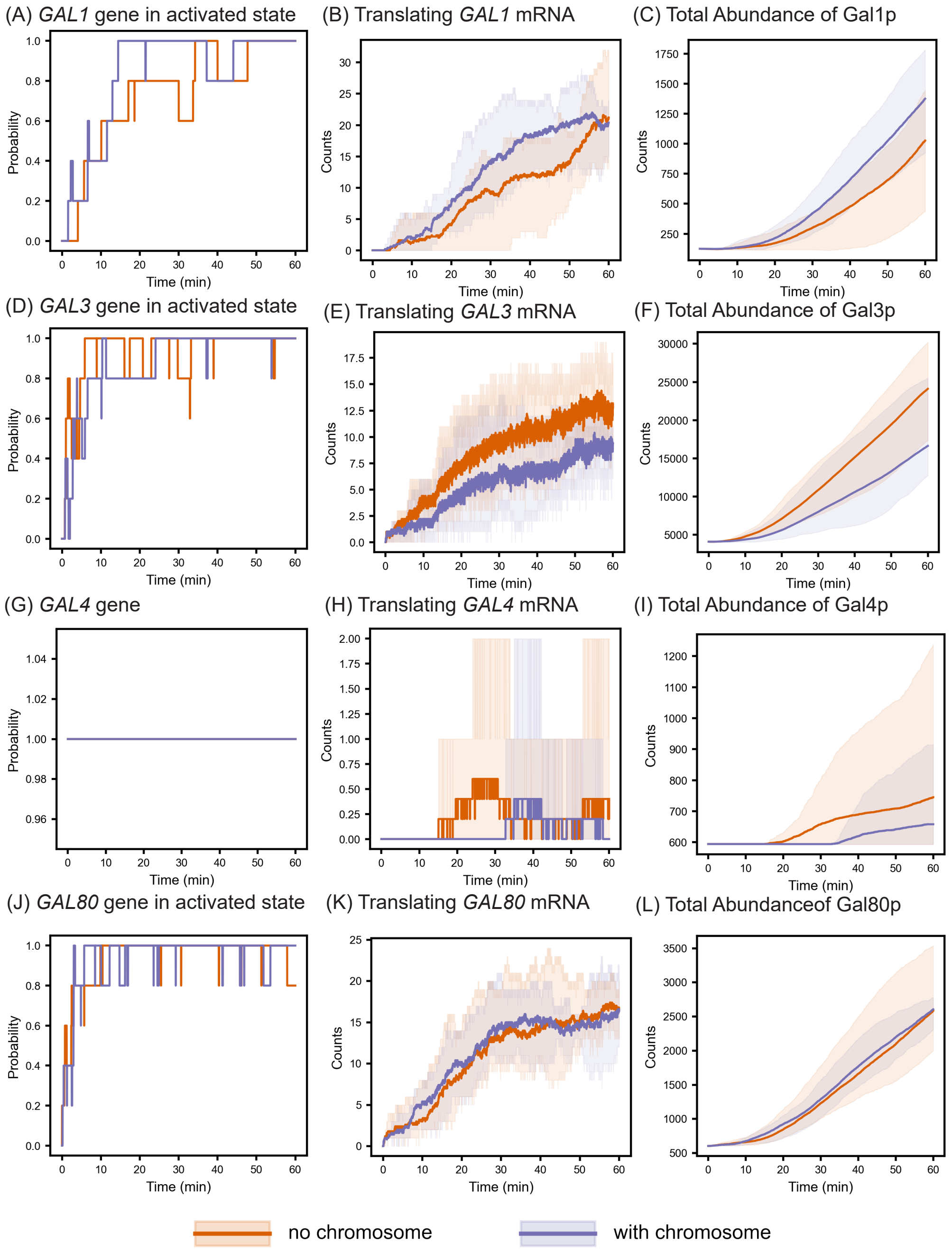

**Figure S4**

#### 3.3 Additional *GAL*-related species for Section 3.3

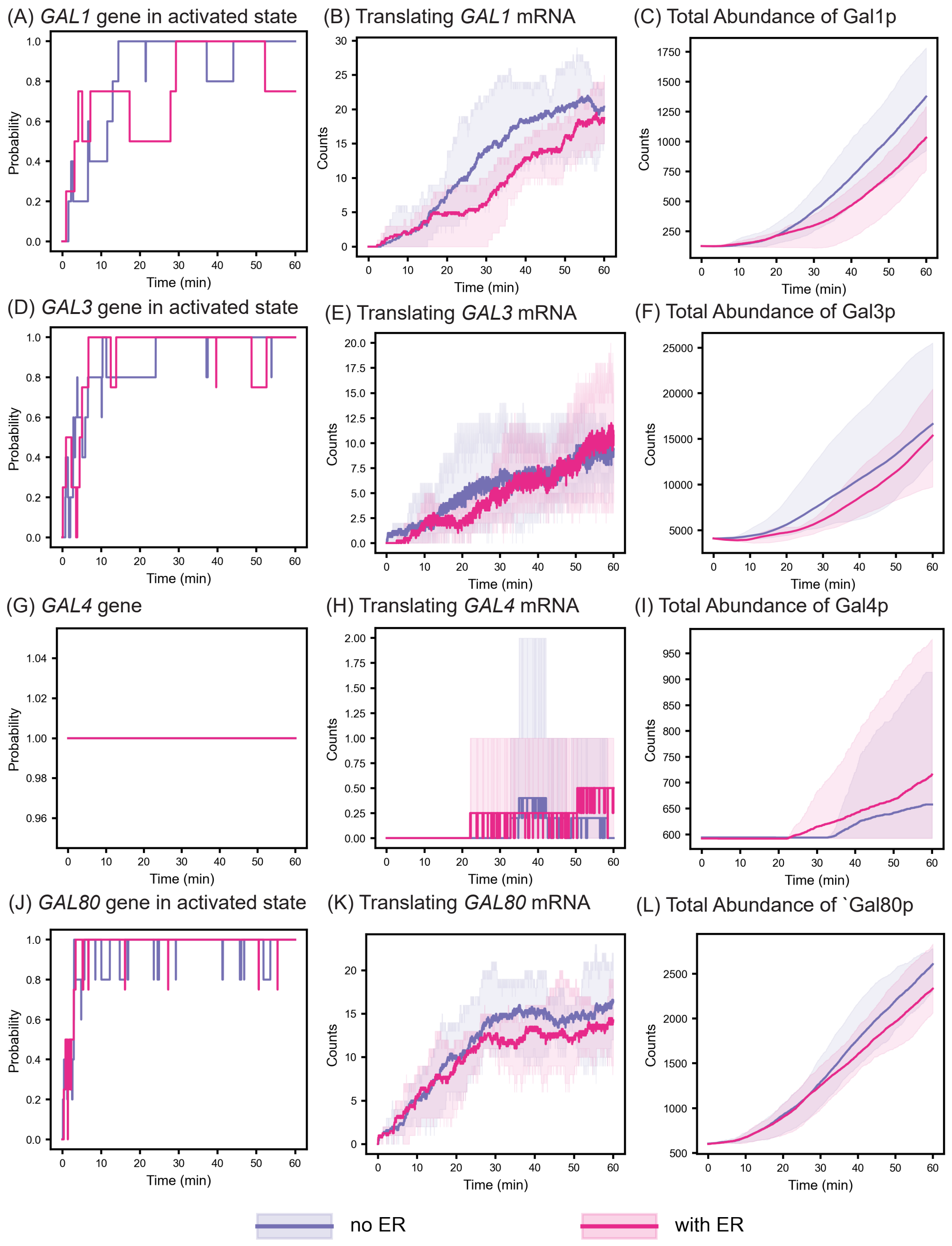

**Figure S5**

#### 3.4 Additional *GAL*-related species for Section 3.4

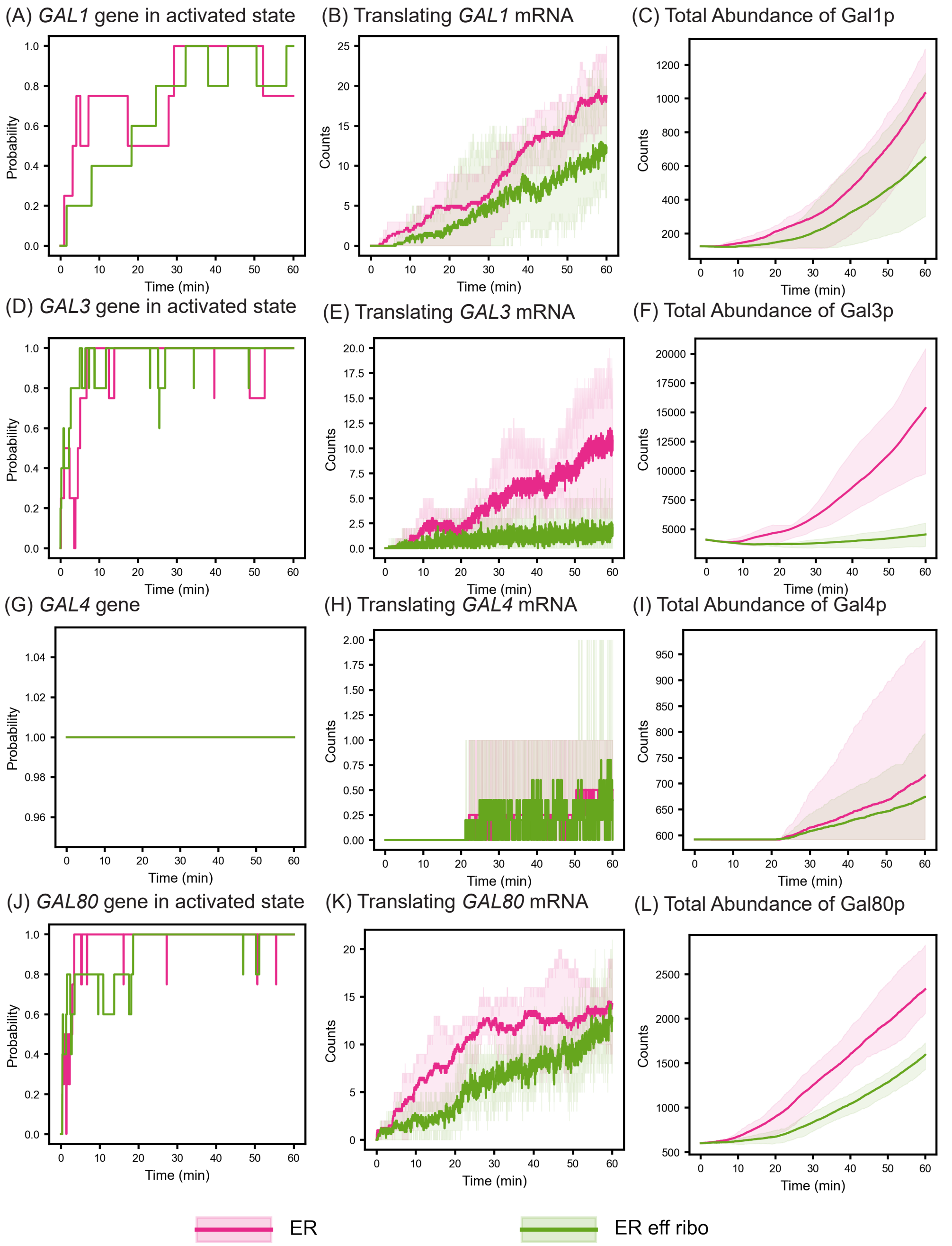

**Figure S6**

As shown in Fig. S6F, the induction effect of Gal3p appears weaker than that of the other *GAL* proteins in the effective-ribosome simulations. This reduced induction primarily arises from the high initial Gal3p abundance used in the model, which was at least fourfold higher than that of the other *GAL* genes. The initial abundance (4,341 molecules) was adopted from Ramsey *et al.* [1] and also applied in the CME study by Bianchi *et al.* [2].

To evaluate the effect of a more physiologically relevant starting level, we estimated the initial Gal3p abundance based on the mean value reported in the Saccharomyces Genome Database (SGD) [6], which is 865 molecules, and scaled it according to proteomics data for cells grown in 2% raffinose medium [7]. Applying the reported ratio (65%), the adjusted initial abundance is $0.65 \times 865 = 564$. When this revised abundance is used, the induction response of Gal3p becomes comparable to that of the other *GAL* proteins, as shown in Fig. S7B.

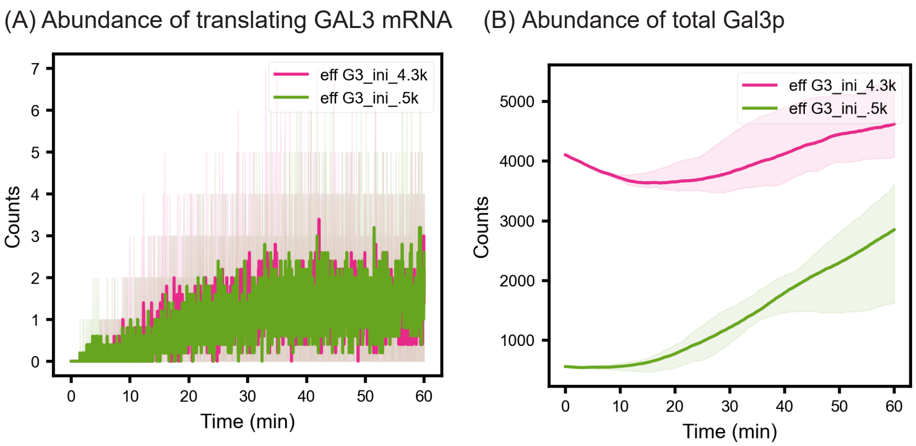

**Figure S7**

At the same time, we note that the initially higher fraction of translating *GAL2* mRNA(Main Text, Fig. 9A) arises from its relatively low total mRNA abundance at the start of the simulation. Moreover, no significant differences were observed within the first 10 minutes in either total *GAL2* mRNA or translating *GAL2* mRNA levels ($p < 0.05$; FigS8).

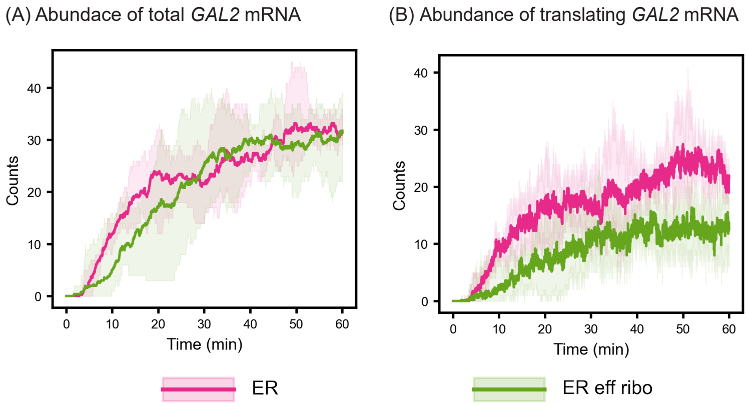

**Figure S8**

#### 3.5 Ribosome in Translation

The translating ribosome trajectories for no-ER, with-ER and no-ER 2,102 ribosome cases respectively. It is clear, the maximum translating ribosomes are below 150 in first 60 minutes simulation in all cases.

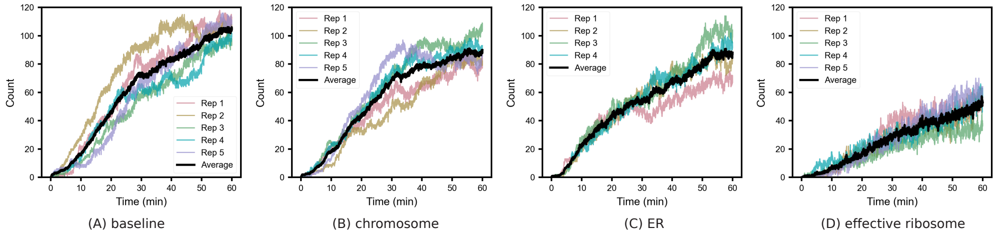

**Figure S9.** (A) Yeast cell in 60 minutes simulation.(B) Yeast cell in 60 minutes simulation with chromosomes only. (C) Yeast cell in 60 minutes simulation with ER and chromosomes. (D) Yeast cell in 60 minutes simulation with effective ribosomes, ER and chromosomes.

#### 3.6 Comparison of Repressor Dynamics in the Nucleoplasm and Cytoplasm Between RDME-ODE and CME-ODE Models

Figure S10 compares the time-resolved dynamics of the repressor G80d in the nucleoplasm (A) and cytoplasm (B) between the well-stirred CME–ODE and spatially resolved RDME–ODE frameworks. In both compartments, the repressor G80d exhibits a rapid transient decline over the first $\sim$10 minutes, followed by relaxation toward a low steady-state abundance for the remainder of the 60-minute simulation window. While the qualitative temporal trend is conserved across modeling frameworks, the spatial RDME–ODE simulations maintain consistently lower repressor copy numbers at early times, most prominently in the nucleoplasm. This early depletion is accompanied by reduced variability in G80d abundance in RDME–ODE compared with CME–ODE, consistent with spatial compartmentalization and diffusion constraints that redistribute repressors across subcellular volumes. Importantly, after the initial transient, both models converge to comparably low nucleoplasmic and cytoplasmic G80d levels, indicating that long-timescale repressor availability becomes similar across frameworks despite differences in early-time spatial redistribution.

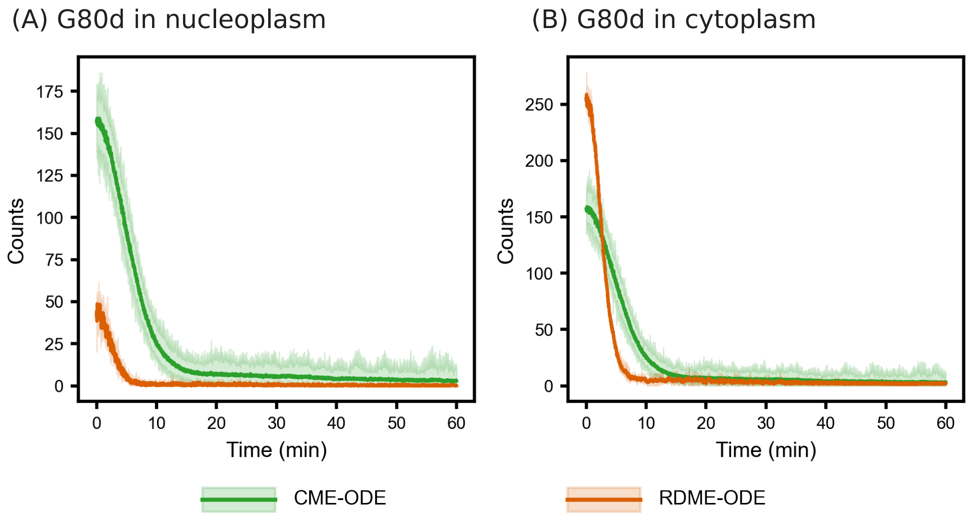

**Figure S10.** (A)G80d abundance in nucleoplasm. (B)G80d abundance in cytoplasm.

##### 3.7 Comparison of Activator and Repressor Dynamics in the Nucleoplasm and Cytoplasm Between Non-Chromosome and Chromosome-Included Models

Figure S11 compares the nucleoplasmic dynamics of the activator Gal4p dimer (G4d; panel A) and the repressor Gal80p dimer (G80d; panel B) between spatial RDME–ODE simulations performed with and without explicit chromosome geometries. Across the full 60-minute simulation window, G4d copy numbers remain relatively stable and exhibit substantial overlap between the two conditions, indicating that inclusion of chromosome occupancy does not measurably perturb the nucleoplasmic abundance of the core activator. Similarly, G80d displays a rapid early-time depletion over the first ∼10 minutes, followed by convergence to near-zero abundance thereafter; importantly, this transient decay profile is nearly identical between chromosome and non-chromosome simulations. Together, these results suggest that introducing chromosome lattice obstacles does not significantly alter the steady-state or transient nucleoplasmic availability of either regulator, consistent with a minimal impact of chromosome volume exclusion on regulator diffusion and promoter accessibility at the occupied volume fractions present in the model.

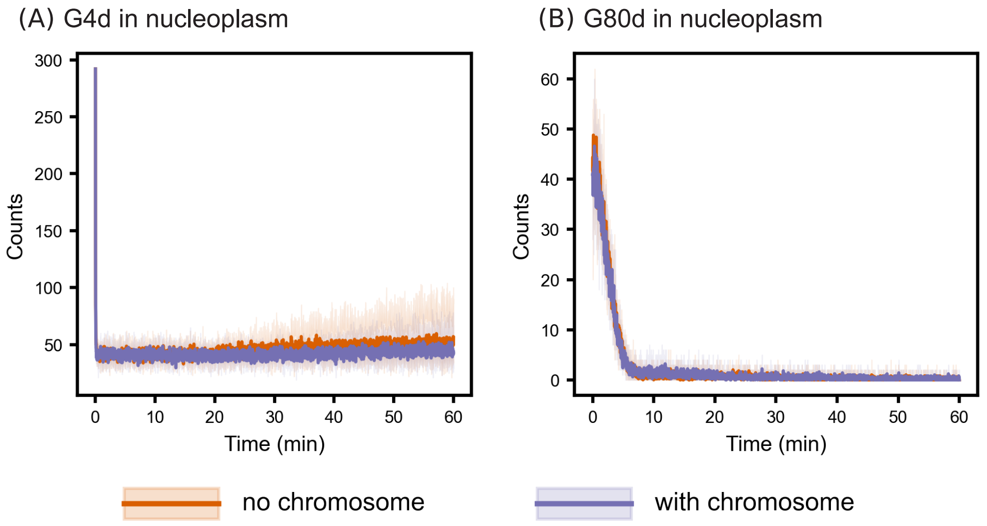

**Figure S11.** (A)G80d abundance in nucleoplasm. (B)G80d abundance in cytoplasm.

##### 3.8 Gene Positioning in the Nucleus Has Minimal Impact on Activation Dynamics

We next examined whether the spatial placement of *GAL* genes within the nucleus can influence induction dynamics in the RDME–ODE framework. To isolate gene-position effects from chromosome volume exclusion, we performed additional simulations using the RDME–ODE model with all chromosome lattice sites removed, yielding a homogeneous nucleoplasmic volume. Within this non-chromosome setting, we tested two extreme gene localization scenarios: (i) all *GAL* network genes placed at the geometric center of the nucleoplasm and (ii) all *GAL* genes placed near the nuclear periphery at randomly selected edge voxels. Across these conditions, we observed no significant differences ($p > 0.05$) in either the timing or magnitude of gene activation, indicating that gene positioning alone does not measurably affect induction under the present parameter regime. For clarity, the plots show only transporter-related species.

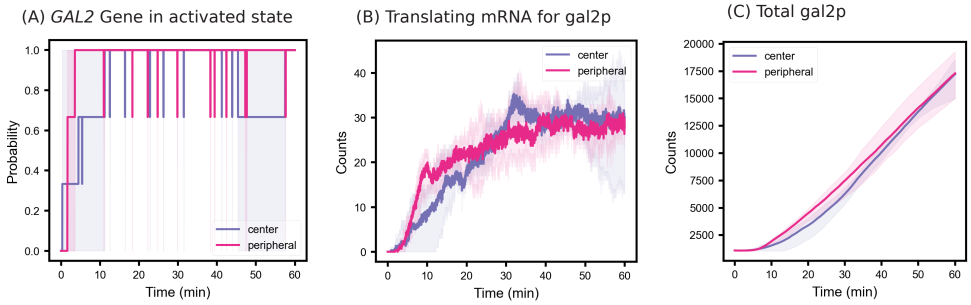

**Figure S12.** RDME-ODE simulations with gene at the center and the peripheral of nucleoplasm.

##### 3.9 Ribosome Number and Spatial Organization Jointly Limit GAL2 Translation

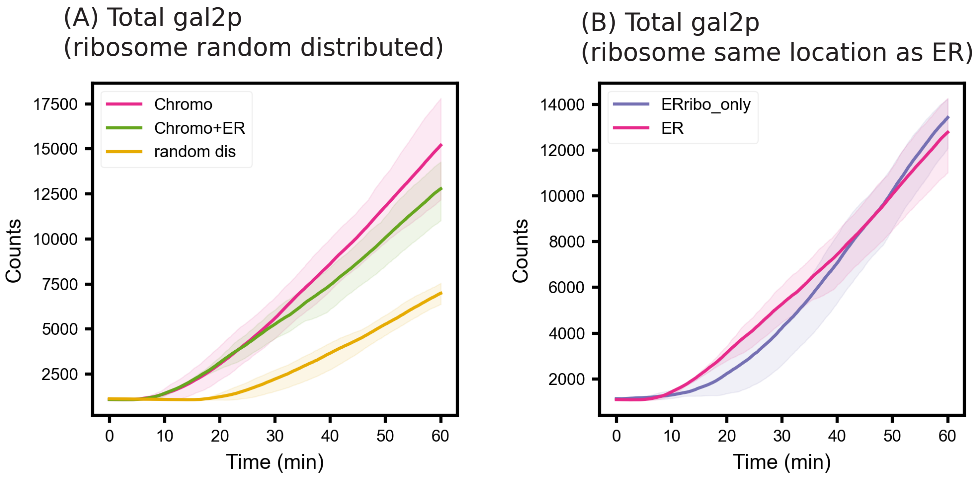

**Figure S13.** Numerical and spatial ribosome distributions jointly shape *GAL2* expression dynamics.(A) Comparison of total gal2p abundance in chromosome (no ER), ER (with ER) and randomly distributed simulation. (B) Comparison of total gal2p in ERRibo_only (ER case with ER geometry removed and keep ER-associated ribosomes in their location) and ER (”with ER” case in main text).

#### 4 Comparison of Predicted and Measured Final Fold Changes of GAL Proteins Across Experimental and Modeling Studies

Both results are close to what Ramsey et al. reported around x10 fold changes for protein Gal1p/Grep in 420 min simulation with Ramsey's original ODE model. [1].

**Table S16.** Fold-change comparison of GAL proteins under two galactose concentrations.

| Species (Total) | 5.55 mM/ 0.1% (Ramsey, ODE) | 11.1 mM/ 0.2 %(Ramsey, ODE) |
|---|---|---|
| Gal1p/Grep | 9.23 | 10.49 |
| Gal2p | 12.81 | 14.44 |
| Gal3p | 6.00 | 6.19 |
| Gal80p | 2.51 | 2.58 |

**Table S17. Comparison of Initial Protein Copy Numbers from Ramsey's [1] and Paulo's Datasets [7].** Values from Ramsey et al. were extracted from the Supplemental Information of the original publication. Values from Paulo et al. represent mean proteomics abundances measured in 2% raffinose medium. All quantities were rounded to the nearest integer to satisfy the requirement that molecular counts be integers in stochastic simulations.

| Species (Total) | Ramsey | Paulo |
|---|---|---|
| Gal1p | 132 | 7733 |
| Gal2p | 1157 | 2184 |
| Gal3p | 4341 | 1777 |
| Gal80p | 3 | 62 |
| Gal4p | 618 | 2033 |
| Reporter protein | 628 | NA |

### 5 Comparison of propensity of second order reaction in CME and RDME

Assuming we have second order reaction as:

$$A + B \longrightarrow C$$

with forward kinetic rate $k$ $(s^-1 \cdot M^-1)$. The system(cell) has volume $V_{sys}$ and Avogadro Number is $N_A$.

For CME, the propensity is clear:

$$P_{CME} = \frac{k}{N_A \cdot V_{sys}} \cdot S_A S_B$$

where $S_A, S_B$ is the total abundance of A and B in the cell.

FOR RDME, in each subvoxel $i$ with volume $V_i$, we have:

$$P_i = \frac{k}{N_A \cdot V_i} \cdot S_{A,i} S_{B,i}$$

where $S_{A,i} S_{B,i}$ is the abundance of A and B in the subvoxel $i$.

Since the subvoxels all have the same volume, we have total number of N subvoxels in 3D lattice:

$$N = V_{sys}/V_i$$

The total propensity for the overall system is:

$$P_{RDME} = \sum_i^N P_i = \sum_i^N \frac{k}{N_A \cdot V_i} \cdot S_{A,i} S_{B,i} = N \cdot \frac{k}{N_A \cdot V_{sys}} \sum_i^N S_{A,i} S_{B,i}$$

In our gene activation model, only one subvoxel $j$ will have gene species, therefore we have one of the species abundance(e.g. A) $S_{A,i} \equiv 0$ if $i \neq j$ and $S_{A,j} = 1 = S_A$. Then, the equation for gene activation/repression can be simplified into:

$$P_{RDME} = N \cdot \frac{k}{N_A \cdot V_{sys}} S_{A,j} S_{B,j} = N \cdot \frac{k}{N_A \cdot V_{sys}} \cdot S_{B,j}$$

Similarly, for CME:

$$P_{CME} = \frac{k}{N_A \cdot V_{sys}} \cdot S_B$$

The ratio of the propensities is:

$$\frac{P_{RDME}}{P_{CME}} = \frac{N \cdot \frac{k}{N_A \cdot V_{sys}} \cdot S_{B,j}}{\frac{k}{N_A \cdot V_{sys}} \cdot S_B} = N \cdot \frac{S_{B,j}}{S_B}.$$

. In the simulations performed in the paper, the total subvoxel number $N = 192^3 >> \frac{S_{B,j}}{S_B} \approx \frac{1}{300}$, and then so the propensity in RDME case is much larger. Similar calculation also holds for gene repression.

### 6 Mathematical interpretation of ER-dependent translation and trafficking

To formalize the impact of endoplasmic reticulum (ER) geometry on transporter expression, we describe the process using a minimal diffusion–encounter and compartmental trafficking framework. Translation initiation of *GAL2* mRNA is treated as a two-step process consisting of (i) diffusion-limited encounter with ER-bound ribosomes and (ii) a local biochemical initiation step. The effective ER-dependent initiation rate is written as

$$\frac{1}{k_{\text{init}}^{\text{ER}}} = \frac{1}{k_{\text{diff}}^{\text{ER}}} + \frac{1}{k_{\text{chem}}}, \qquad k_{\text{diff}}^{\text{ER}} \sim 4\pi D_{\text{mRNA}} a \rho_{\text{ER-ribo}},$$

where $D_{\text{mRNA}}$ is the effective diffusion coefficient of *GAL2* transcripts, $a$ is an effective capture radius, and $\rho_{\text{ER-ribo}}$ is the surface density of ER-bound ribosomes.

Following synthesis, Gal2p dynamics are modeled using a two-compartment trafficking system:

$$\frac{dG_{\text{ER}}}{dt} = J_{\text{syn}}(t) - k_{\text{export}} G_{\text{ER}} - k_{\text{deg,ER}} G_{\text{ER}},$$

$$\frac{dG_{\text{PM}}}{dt} = k_{\text{export}} G_{\text{ER}} - k_{\text{int}} G_{\text{PM}} - k_{\text{deg,PM}} G_{\text{PM}},$$

where $G_{\text{ER}}$ and $G_{\text{PM}}$ denote Gal2p abundance in the ER and plasma membrane compartments, respectively.

ER geometry enters the model through an effective trafficking time,

$$\tau_{\text{ER}}^{\text{geom}} \sim \frac{\left(L_{\text{ER}}^{\text{eff}}\right)^2}{2D_{\text{ER}}} + \frac{L_{\text{ER}}^{\text{eff}}}{v}, \qquad k_{\text{export}} \approx \frac{1}{\tau_{\text{ER}}^{\text{geom}}},$$

where $L_{\text{ER}}^{\text{eff}}$ is an effective path length within the ER network, $D_{\text{ER}}$ is the lateral diffusion coefficient of Gal2p, and $v$ represents any directed transport component. Increased ER geometric complexity therefore decreases $k_{\text{export}}$, lowering steady-state plasma-membrane abundance

$$G^*_{\mathrm{PM}} \approx \frac{k_{\mathrm{export}}}{k_{\mathrm{int}} + k_{\mathrm{deg,PM}}} \cdot \frac{J_{\mathrm{syn}}}{k_{\mathrm{export}} + k_{\mathrm{deg,ER}}}.$$

This reduction in plasma-membrane Gal2p directly limits galactose uptake, thereby reducing intracellular galactose levels.

### 7 Sensitivity Analysis on kinetic parameters

To further support our argument that the increased probability of the GAL2 gene being in the activated state leads to higher levels of G2 protein translation within the RDME–ODE framework, we conducted a parameter sensitivity analysis. Specifically, we first constructed a pure ODE model based on the CME–ODE formulation and systematically varied all kinetic parameters within a range of $0.1\times$ to $10\times$ their baseline values. Parameters that produced more than a twofold change in G2 abundance after 60 minutes of simulation were identified(Fig S14(B)). We then generated pairwise heatmaps of these parameters to visualize their combined effects. Notably, the diffusion rates of G80d between the cytoplasm and nucleoplasm were capable of driving more than a twofold change in G2 abundance together(Fig S14(A, C)). Moreover, when two such parameters were altered simultaneously, the resulting G2 abundance closely approximated that predicted by the RDME–ODE model at 60 minutes(Fig S14(D)). These findings support our conclusion that the observed increase arises primarily from asymmetric diffusion.

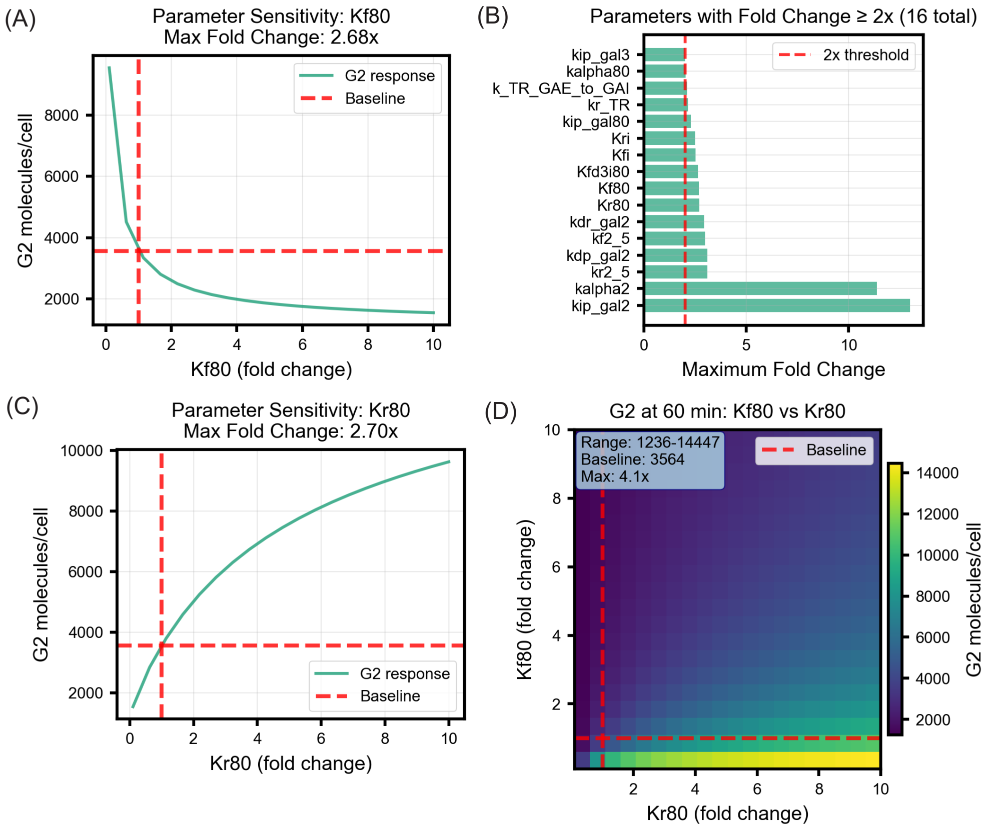

**Figure S14.** Parameter Sensitivity Analysis Results. (A) Effect of single-parameter perturbation of the G80d diffusion rate $kf80$ from the cytoplasm to the nucleoplasm. (B) Kinetic parameters exhibiting more than a twofold change in G2 abundance within the 0.1× to 10× perturbation range. (C) Effect of single-parameter perturbation of the G80d diffusion rate $Kr80$ from the nucleoplasm to the cytoplasm. (D) Combined perturbation of both G80d diffusion rates (cytoplasm → nucleoplasm and nucleoplasm → cytoplasm).
